# Boxcar Imaging FCS Reveals Membrane Raft Stabilization Kinetics in Antigen-Stimulated Mast Cells

**DOI:** 10.64898/2025.12.16.694662

**Authors:** Gil-Suk Yang, Nirmalya Bag, Barbara A. Baird

## Abstract

Antigen (Ag) crosslinking of immunoglobulin E–receptor (IgE-FcεRI) complexes in mast cells and consequent coupling with Lyn tyrosine kinase in the plasma membrane inner leaflet stimulates transmembrane signaling to initiate allergic and inflammatory responses. As established previously, this coupling requires formation of liquid-ordered (Lo)-like regions (aka “rafts”) around the nano-clustered receptors to facilitate lipid-based partitioning of Lyn via its membrane anchor, followed by receptor phosphorylation mediated by protein-protein interactions. Imaging fluorescence correlation spectroscopy (ImFCS) was previously used to measure diffusion of Lyn-EGFP and its lipid anchor PM-EGFP (both Lo-preferring) as well as EGFP-GG (inner leaflet lipid probe, liquid-disordered (Ld)-preferring) and showed that the membrane reorganized within 15 minutes after Ag addition. To quantify the transition kinetics between the resting and Ag-stimulated steady-states, we have now developed Boxcar ImFCS for time-resolved diffusion measurements on sub-minute scale. We found that Ag stimulation causes gradual diffusion decreases for Lyn-EGFP and PM-EGFP with distinctive half-times (*t_1/2_*) of 6.9 min and 12 min, respectively, showing that Lyn’s protein-based interactions accelerate its diffusional transition. Simultaneously, EGFP-GG gradually changes to faster diffusion with *t_1/2_* = 9.4 min. In comparison, *t_1/2_* = 5.0 min for recruitment of cytoplasmic Syk by phosphorylated FcεRI, consistent with initiation of transmembrane signaling before global membrane reorganization and raft condensation is completed by large, stabilized Ag-IgE-FcεRI clusters. Boxcar ImFCS extends the analytical power of ImFCS to reveal dynamic membrane processes that may accompany stimuli-receptor interactions and their sequalae.

**STATEMENT OF SIGNIFICANCE:** Stimulated lipid reorganization and stabilization of liquid-ordered (Lo)- like regions (“rafts”) in the plasma membrane inner leaflet are decisive for initiating IgE-receptor-mediated mast cell signaling. Here, we developed a new technique, termed Boxcar Imaging Fluorescence Correlation Spectroscopy, to determine the kinetics of raft stabilization after antigen binding and crosslinking IgE receptors. We provide one of the first characterizations of time-dependent raft condensation as stimulated in live cells. We envisage broad applications of this experimental strategy to quantitatively decipher intertwined processes of membrane phase-like separation and functional transmembrane signaling.

## INTRODUCTION

Presence in the heterogeneous plasma membrane of ordered-lipid regions characterized by cholesterol and lipids with saturated fatty acids – colloquially known as “rafts” (1–3) – has been established, although their biochemical and biophysical nature and stability remain hotly debated (4–7). The physical properties of these ordered-lipid regions resemble those of a liquid-ordered (Lo) phase, which co-exists with a liquid-disordered (Ld) phase, in synthetic vesicles made of specific lipid composition (8–10), and in cell-derived giant plasma membrane vesicles (11, 12), both at equilibrium. In the plasma membranes of living, resting cells, these Lo-like, ordered-lipid nanodomains are transient in that, on average, they form and disappear every tens of milliseconds (13, 14). This transience likely contributes in large part to observed marginal differences between the physical properties of ordered regions and surrounding Ld-like, disordered regions at the spatio-temporal scales of current high-resolution techniques (13, 15–17). The phase-like organization can be stabilized by collective interactions between membrane constituents that are imposed by external factors such as physical crosslinking of membrane proteins or lipids (2, 18). Compelling evidence supports the view that this phase-like behavior of lipids is important for regulating protein interactions in the membrane, as described herein, and these may also couple to phase-based protein condensates in the cytoplasm (19).

Stimulated transmembrane signaling through cell surface immunoreceptors that are clustered by means of multivalent antigen (Ag) is one of most studied processes where changes in phase-like organization is shown to play a decisive role (20–25). The high affinity receptor for IgE (FcεRI) in rat basophilic leukemia (RBL) mast cells has provided a robust experimental system for examining this process. For example, our recent imaging fluorescence correlation spectroscopy (ImFCS) measurements clearly demonstrated stabilization of ordered regions in the inner leaflet of RBL plasma membranes in the Ag-stimulated steady-state (22) (Fig. 1). We utilized genetically encoded, inner leaflet lipid probes, namely enhanced green fluorescent protein (EGFP)-labeled palmitoylated/myristoylated-peptide (PM-EGFP) and geranyl-geranylated-peptide (EGFP-GG) (Fig. 1; (26)), and we measured average diffusion coefficients (*D_av_*) at very high precision using ImFCS (standard error ±1%) in both resting (before Ag-crosslinking) and stimulated (15 min after Ag-crosslinking of IgE-FcεRI) steady-states (Fig 1; (22)). PM-EGFP prefers ordered regions of the membrane enriched in saturated lipids, while EGFP-GG prefers disordered regions enriched in unsaturated lipids (26). Since these non-functional inner leaflet probes have no protein-interacting components, their diffusion behavior is determined primarily by lipid-based interactions. We observed that with Ag-crosslinking of IgE-FcεRI the *D_av_* of PM-EGFP decreases 8% in stimulated steady-state while *D_av_* of EGFP-GG increases by 10%. These results are consistent with net stabilization of ordered and disordered regions in the Ag-stimulated steady-state. Our results thus demonstrate plasma membrane ‘adaptability’(27) from transient to more stable phase-like organization after Ag-clustering of IgE-FcεRI. We further showed that the *D_av_* of Lyn-EGFP, which is order-preferring and contains the same peptide segment and lipid anchor as PM-EGFP (Fig. 1; (26)), decreases by 10% in stimulated steady-state while that of a disorder-preferring Lyn mutant, S15-Lyn-EGFP, increases by 4%. This distinctive behavior reflects the coupling of Lyn-EGFP, but not of S15-Lyn-EGFP, with Ag-clustered IgE-FcεRI: Lyn-EGFP, but not S15-Lyn-EGFP, was shown to facilitate Ag-dependent phosphorylation in a reconstituted system. This means that the lipid-based partitioning preference of Lyn is key to its functional coupling as mediated by stabilization of ordered regions around Ag-crosslinked IgE-FcεRI (20). Lyn’s protein domains, SH2, SH3 and kinase, then allow direct binding and phosphorylation of FcεRI, followed by recruitment of cytoplasmic Syk tyrosine kinase (28). In another study on the same Ag-stimulated signaling system, we showed that cells with more ordered resting plasma membrane produce stronger stimulated responses, including Syk recruitment, and the opposite effect when the plasma membrane is less-ordered (29). More recently, examination with microvillar cartography showed that IgE-FcεRI, Lyn, and ordered-lipid nanodomains localize to membrane microvilli in suspended resting RBL cells, where they may serve as signaling hubs upon addition of Ag (30). Taken together, our results over multiple studies provide compelling evidence for functional competency of ordered regions in the plasma membrane to initiate IgE-FcεRI signalling (31).

**Figure 1:**
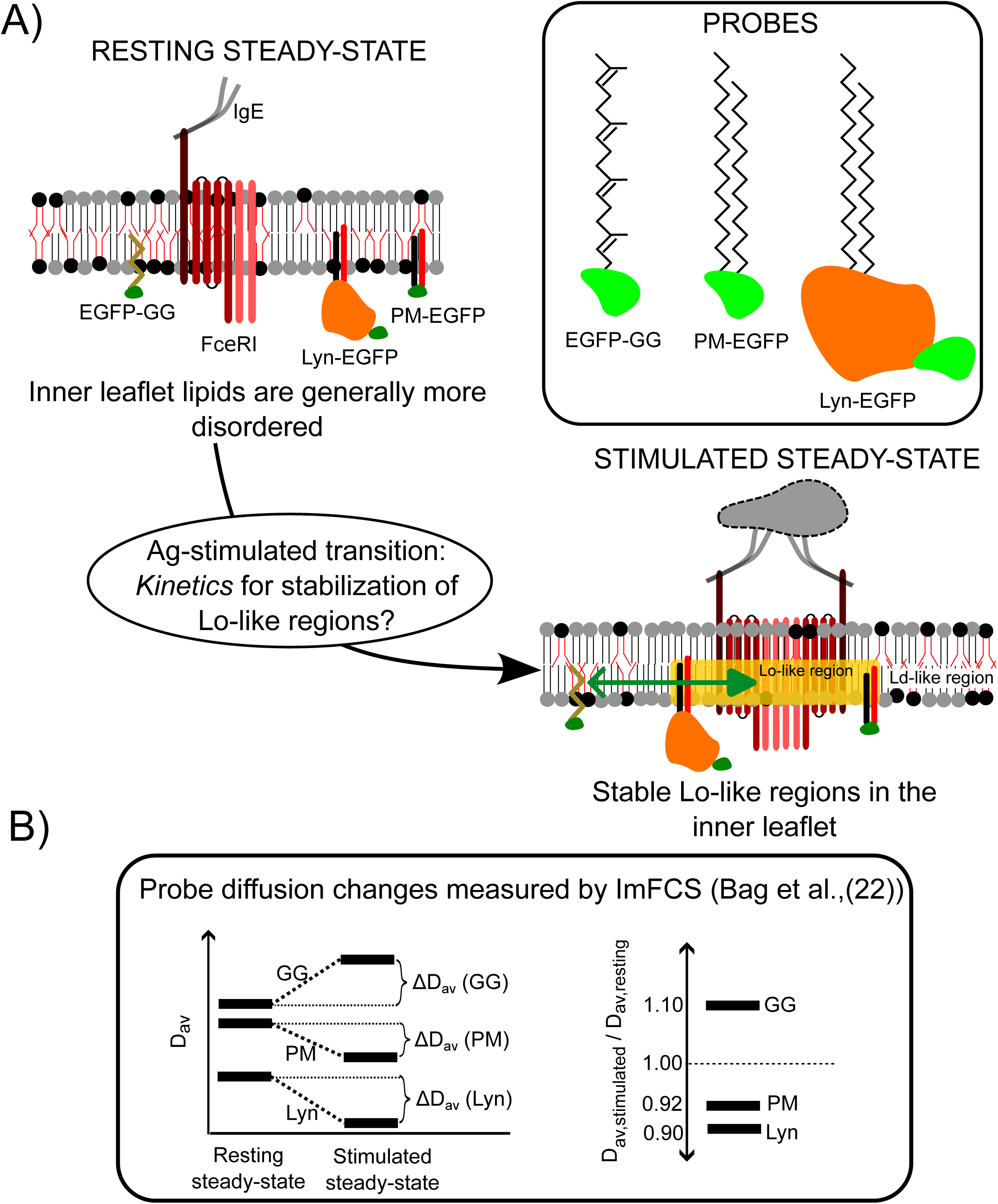
Antigen-mediated clustering of IgE-FcεRI initiates transmembrane signaling and reorganization of plasma membrane components as can be measured with changes in probe diffusion. A) Schematic of the experimental system including IgE-FcεRI, antigen (Ag), and membrane probes. Ag-stimulated changes in membrane organization to stabilize Lo-like regions are measured by changes in average diffusion coefficient (*D_av_*) of probes. Structures in schematic are not drawn to scale: IgE (M_r_∼200,000); FcεRI ( M_r_∼70,000), Lyn (M_r_∼60,000), lipid tails (M_r_<1,000), and EGFP (M_r_∼30,000). B) Changes in steady-state *D_av_* of specified probes before and after 15 min of Ag stimulation, as measured previously with ImFCS (22). The present study focuses on the transition kinetics of membrane re-organization and stabilization of Lo- and Ld-like membrane domains that occur in the inner leaflet after Ag stimulation.

The observations summarized above show that the transient ordered nanodomains in the plasma membranes of resting cells grow into stable and signaling-competent platforms after IgE-FcεRI crosslinking by Ag. Since cellular signaling events are tightly controlled in space and time (32), it is important to elucidate how the plasma membrane responds to Ag and other potential stimuli to stabilize phase-like organization that initiates transmembrane signaling and possibly also facilitates downstream processes. Our present study aimed to evaluate the time dependence of inner leaflet ordering and the mode (abrupt *vs* gradual) of transition between resting and stimulated membrane steady-states by monitoring diffusion of order-preferring (PM-EGFP, Lyn-EGFP) and disorder-preferring (EGFP-GG) probes. For this purpose, we developed an analysis scheme for time-resolved diffusion changes, termed Boxcar ImFCS, to determine *D_av_* of these membrane components at sub-minute intervals. Our measurements monitor the membrane’s biophysical response after addition of Ag when phase-like features become globally stabilized. We compare the time-dependence of stimulated changes in probe diffusion to that of stimulated recruitment to the plasma membrane of cytoplasmic YFP-Syk. We show ordered regions continue to grow as Ag crosslinking of FcεRI into larger aggregates continues until a steady-state is reached. This level of stabilization represents an amplification of phase-like changes occurring with the nanoscale aggregates that are sufficient to initiate FcεRI-Lyn coupling and thereby transmembrane signaling (20). Our work substantially extends the capabilities of ImFCS to measure time-dependent changes in probe diffusion with sub-minute resolution. This novel measurement and analysis scheme is expected to be generally applicable to a broad range of membrane biophysics questions.

## MATERIALS AND METHODS

Detailed Materials and Methods are described in the Supplemental Information and Results sections. Briefly, RBL-2H3 cells in culture were chemically transfected with a specified plasmid using FUGENE. ImFCS measurements were performed on the transfected cells covered with buffered saline solution in a 35-mm glass-bottomed dish mounted on a home-built TIRFM system attached with a fast EMCCD camera. ImFCS measurements were done on a region of interest (ROI) on the verntral surface of a cell before addition of antigen (-Ag, resting steady-state condition) and after Ag addition (+Ag condition). The time between Ag addition and the start of ImFCS measurements are described in the Results section. All measurements were done at room temperature. The raw data were analyzed with ImFCS plug-in (33) for ImageJ/FIJI (34) (for determination of diffusion coefficient map) and Igor Pro 8.0 (WaveMetrics, OR; for bootstrapping and statistical tests). Simulations were done in Wolfram Mathematica and Matlab (The MathWorks, Inc., MA).

## RESULTS

We developed Boxcar ImFCS to evaluate for selected probes the kinetics of diffusional transition from resting to stimulated steady-states that accompanies transmembrane signaling initiated by Ag-crosslinking of IgE-FcεRI in RBL mast cells. As demonstrated in our previous work (20, 22, 24), particular protein-based and lipid-based interactions underlie the coupling of Ag-crosslinked FcεRI and Lyn kinase. This coupling requires stabilization of Lo-like, ordered-lipid regions around the clustered receptors, facilitating lipid-based partitioning of Lyn via its membrane anchor. This partitioning serves to increase the frequency of the protein-based interactions (via Lyn’s SH2, SH3 and kinase domains) leading to effective phosphorylation of FcεRI cytoplasmic segments. Since the membrane protein and lipid components undergo selective interactions, it is likely that the nature of their transition from resting to stimulated steady-states will be different. Lyn-EGFP is involved in both lipid-based and protein-based interactions whereas PM-EGFP (the lipid-anchor of Lyn) only participates in lipid-based interactions. Correspondingly, Boxcar ImFCS analyses of PM-EGFP can measure the formation of stable Lo-like regions in the inner leaflet, which are sensed by saturated lipids. The rate of this process, when compared to that measured for Lyn-EGFP would thus reflect how additional protein-based interactions modulates the transition rate. For further comparison, measured changes in EGFP-GG diffusion reflect the formation of Ld-like, disordeded-lipid regions after Ag-stimulation. Thus, we sought to evaluate each of these three probes at sequential time points after Ag addition until their diffusion properties reached the stimulated steady-state.

### Part I: Development of Boxcar ImFCS to detect subtle transition between plasma membrane steady-states

#### Large ImFCS data sets can be segmented to enable time resolution of successive diffusion measurements

Since its inception, FCS (35, 36) and its camera-based modality, ImFCS (37), have been used to quantify the steady-state diffusion of probes in biophysical systems (38, 39). As illustrated in Fig. S1A-C, a typical ImFCS measurement is conducted on the ventral surface of an adherent cell, within a region of interest (ROI) consisting of 25×25 Px units (1 Px unit = 320×320 nm^2^ in our set-up) and takes 280 sec comprising 80,000 movie frames (*n_f_* = 80,000) recorded at 3.5 msec per frame (15). Operative terms are defined in Table 1. Individual fittings of all 625 autocorrelation functions (ACFs) from these raw data yields a diffusion coefficient (*D*) value of the selected probe for each spatially identified Px unit. This set of *D* values can be presented as a spatial map of diffusion coefficients (*D* map) or as normalized *D* distributions, including probability distribution functions (PDF) and cumulative distribution functions (CDF) (Fig. S1D-F) (15, 22, 40). The *D* map provides insight into spatial heterogeneities that affect probe diffusion in individual Px units, such as spatially restricted diffusion obstacles imposed by bio-active peptides (41, 42). The *D* distributions yield statistical information that can be subjected to further analysis of diffusion properties (15). Pooling all Px unit data from 15-20 cells gives ∼10,000 *D* values, from which an average diffusion coefficient (*D_av_*) can be determined. Calculated in this manner, the standard error of the *D_av_* values for a variety of membrane probes under a range of conditions are typically less than 1% (15, 22, 40). Diffusion measurements of lipid and protein probes with such high precision before (i.e., resting steady-state) and after a treatment (i.e., perturbed steady-state) have been instrumental in delineating subtle changes in membrane organization caused by ligand engagement with specific receptors (22, 40, 43), cholesterol extraction (40), inhibition of actin polymerization (15, 40), and transbilayer interactions (44, 45, 29).

**Table 1:** Term Definitions.

| Symbol | Parameter |
| --- | --- |
| $D$ | Diffusion coefficient |
| $D_{av}$ | Average diffusion coefficient of an ROI |
| $D_0$ | Average diffusion coefficient in resting steady-state condition |
| $D_t$ | Average diffusion coefficient at time $t_{stim}$ after Ag addition |
| Px unit | Smallest size of membrane unit where $D$ is measured |
| $n_f$ | Total number of frames used for boxcar ImFCS analysis, typically 80,000 in this study |
| $n_{boxcar}$ | Number of frames per boxcar used for ImFCS analysis, typically 40,000 in this study |
| $\Delta n_{boxcar}$ | Number of frames shifted for consecutive boxcar ImFCS analysis: $\Delta n_{boxcar} = n_{boxcar}$ for non-overlapping boxcars; $\Delta n_{boxcar} < n_{boxcar}$ for overlapping boxcars, typically 10,000 in this study |
| $t_{boxcar}$ | Midpoint time of a specified boxcar from initiation of the measurement |
| $\Delta t_{boxcar}$ | Time shift between consecutive boxcar ImFCS analysis: $\Delta t_{boxcar} = t_{boxcar}$ for non-overlapping boxcars; $\Delta t_{boxcar} < t_{boxcar}$ for overlapping boxcar ImFCS analysis, typically 35 sec in this study |
| $t_{stim}$ | Stimulation time after Ag addition |
| $t_{on}$ | $t_{stim}$ at which stimulated change of $D_{av}$ is detected by boxcar ImFCS measurements (10% of total change as fit by Eq. 1) |

In a steady-state, the *D*_av_ value is expected to remain the same over time. As an example, the resting steady-state diffusion of inner-leaflet lipid probe PM-EGFP (Fig. 1) is demonstrated as follows (Fig. 2). We first divide the raw image stack of *n_f_* = 80,000 from an ROI of 25×25 Px units into two non-overlapping segments (boxcars), each comprising 40,000 frames (*n_boxcar_* = 40,000 corresponding to 140 sec acquisition time) (Fig. 2A), and we perform ImFCS analysis on each boxcar to obtain *D* values for each Px unit (Fig. 2B). The spatial map of *D* values represents a picture of all nanoscale phenomena that affect probe diffusion within each Px unit, as averaged over the set time interval. The *D* values of individual Px units in the *D* maps shown in Fig. 2B generally fluctuate over time, while CDFs of all *D* values remain the same (Fig. 2C,D). This is expected for a dynamic system in a steady-state. A closer look at the *D* maps reveal some variations of these natural fluctuations. For example, in Fig. 2B the Px unit indicated by right arrow changes from faster to slower *D* value, whereas the Px unit indicated by left arrow exhibits a relatively low *D* value for both 140-sec time intervals. The latter case suggests the localized presence of relatively stable membrane components that hinder PM-EGFP’s diffusion (15). Thus, these D maps are useful for identifying location and stability of membrane regions exhibiting restricted diffusion under specified conditions and time intervals.

**Figure 2:**
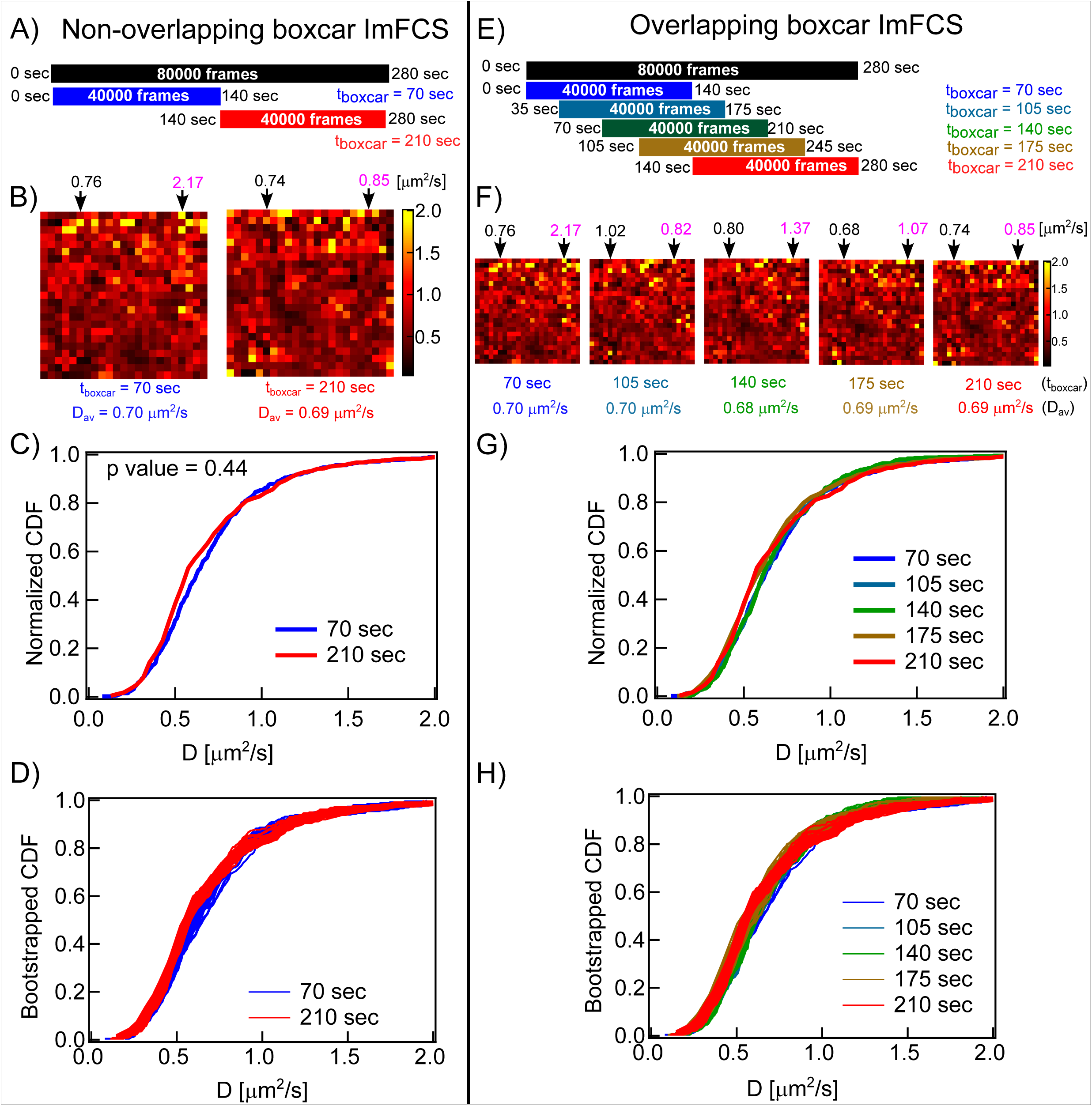
Boxcar ImFCS breaks down image stack into time segments to enable kinetic analysis. Demonstration of Boxcar ImFCS analysis with representative measurement on cell expressing PM-EGFP. <u>Non-overlapping Boxcars</u>: A) An image stack (*n_f_* = 80,000 frames, black bar) is segmented into two non-overlapping boxcars of equal length, i.e, *n_boxar_* = 40,000 and Δ*n_boxar_* = 40,000 (blue and red bars). The actual measurement time range (i.e., 0-140 sec and 140-280 sec) and the corresponding midpoint, *t*_boxcar_ for both boxcars are specified. B) Spatial maps of *D* corresponding to *t*_boxcar_ of 70 sec and 210 sec obtained from ImFCS analyses of individual Px units (see Fig. S1D). The *Dav* values which average over the entire map are given below the maps. Arrows indicate representative individual Px units with *D* values that remain similar (left arrow) or change markedly (right arrow). The *D* values (in μm^2^/s) are given above the representative Px unit. C-D) Obtained from each boxcar are normalized CDF of all *D* values (C) and 30 randomly bootstrapped CDFs using 50% of all *D* values (D). <u>Overlapping boxcars</u>: E) The same image stack as for (A) is segmented into overlapping boxcars of equal length, i.e., *n_boxar_* = 40,000 but with Δ*n_boxar_* = 10,000, as shown by differently colored bars. The resulting ImFCS analyses are assigned to midpoints *t*_boxcar_ as specified F) Spatial maps, G) CDFs of all *D* values (range of *p*-values for pairs of *D* distribution is 0.32-0.89), and H) 30 bootstrapped CDFs using 50% of all *D* values for each of overlapping boxcars. Arrows in spatial maps of (F) point to the same two Px units as in (B), which exhibit similar (left arrow) and different (right arrow) *D* values.

The *D*_av_ value for the first boxcar (1 – 40,000 frames) represents the averaged diffusion coefficients between 0-140 sec from the initiation of measurement, with midpoint = *t_boxcar,1_* = 70 sec. We assign this *D*_av_ value (and the respective *D* map and *D* distribution) to this midpoint.

Similarly, ImFCS analyses on the second boxcar of same size (*n*_boxcar_ = 40,000) but shifted by 40,000 frames (Δ*n*_boxcar_ = 40,000) measures diffusion during frames 40,001 – 80,000 (i.e., between 140-280 sec; midpoint = *t*_boxcar,2_ = 210 sec). The time interval between the midpoints of these two non-overlapping boxcars (Δ*t*_boxcar_) is 140 sec. The principles of boxcar segmentation of image stack were used previously in the context of post-processing of optical imaging such as photobleaching correction (46). For the resting steady-state condition *D_av_* over all 625 Px units at *t*_boxcar,1_ = 70 sec and at *t*_boxcar,2_ = 210 sec are very close (0.7 μm^2^/s), as expected, and the calculated p-value = 0.44 further supports no significant difference (Fig. 2C).

For further evaluation, we compared the bootstrapped CDFs of the *D* values from each boxcar (Fig. 2D). We previously introduced and rigorously characterized bootstrapping of *D* values to reliably delineate small differences of probe’s diffusion between two conditions (22). Bootstrapping analysis is used to mitigate some issues related to conventional hypothesis testing to determine statistical significance of differences between two conditions having large number of data points, as for our ImFCS measurements. In general, p-values decrease exponentially with sample size (47), and statistical tests between two distributions having large sample size often return a very small p-value. Our bootstrapping employed subsampling of the raw distributions (randomly sampling 50% of all data for each distribution, i.e., ∼5,000 *D* values) and then tested the statistical differences pair-wise between bootstrapped subsamples in two independent sets (e.g., two non-overlapping boxcars in resting condition -- or measurements before and after Ag-stimulation, as described later). We repeat this process 30 times resulting in a range of pair-wise p-values that then can be used to evaluate the level of statistical difference. More simply, visual inspection of bootstrapped curves for two independent sets of measurements qualitatively indicates whether these two sets do or do not overlay to exhibit a statistically significant difference.

The bootstrapped *D* CDFs obtained from the two non-overlapping boxcars that are sequential within the same total measurement overlay each other (Fig 2D), confirming overall indistinguishability between diffusion properties measured in the respective 140 sec intervals of the resting steady-state. Unchanged diffusion properties for other inner leaflet probes (EGFP-GG and Lyn-EGFP), outer leaflet probe (YFP-GL-GPI), and transmembrane probe (YFP-GL-GT46) (Fig. S2) confirms the steady-state of resting cells in our study. Similar evaluations of cell membranes at resting steady-state and of supported lipid bilayers have been reported (48).

#### Overlapping boxcars markedly improves the time resolution for successive ImFCS analyses within the same raw data

The time interval between non-overlapping boxcar ImFCS analyses is 140 sec (Fig. 2A) and each of the corresponding *D_av_* values is thereby obtained from a t ∼2.5 min measurement. Since the stimulated steady state previously determined for these probes is established 15 min after Ag addition (22), only six time-dependent *D_av_* values (*D_t_*) could be obtained for this entire time range using a sequential series of four 80,000 frame movies and a 40,000 frame non-overlapping image segmentation strategy. However, we have determined that six data points for *D_av_*vs stimulation time plot are not sufficient to define the transition. Therefore we considered an overlapping boxcar strategy. We first determined the optimal value for *n*_boxcar_. As established previously for bright organic fluorophores as probes in supported lipid bilayers, 20,000 frames is minimally required in ImFCS analysis for accurate and precise determination of *D_av_* (49, 50). More frames are required for probes that diffuse slowly.

Generally, the quality of ACFs improves with the number of frames evaluated and consequently the precision of *D*_av_. However, a larger number for *n_boxcar_* increases the number of images averaged which may increase the “drag” effect during a *D*_av_ transition (see Discussion). To establish the minimum number of frames required for sufficiently precise ImFCS evaluation of time course we used the process illustrated in Fig. S3A, and we confirmed *n*_boxcar_ = 40,000 as optimal for our analyses.

As demonstrated with the resting steady-state in Fig. 2E, decreasing the time interval between two successive ImFCS analyses can be done by segmenting the image stacks into overlapping boxcars. The first boxcar consists of 40,000 image frames (0-140 sec; *n*_boxcar_ = 40,000; 1-40,000 frames of the recorded image stack of 80,000 frames). ImFCS analysis determines *D*_av_ corresponding to *t*_boxcar,1_ = 70 sec (midpoint). For the second boxcar, we shift 10,000 frames (Δ*n*_boxcar_ = 10,000) maintaining *n*_boxcar_ = 40,000 (i.e., 10001-50000 frames; 35-175 sec) and determine *D*_av_ for this time interval, assigned to *t*_boxcar,2_ = 105 sec (midpoint). In this manner, the time interval between two overlapping boxcars (Δ*t*_boxcar_) decreases to 35 sec compared to 140 sec for non-overlapping boxcars. The process for overlapping boxcars can be repeated with Δ*n*_boxcar_ = 10,000 and *n*_boxcar_ = 40,000 for an image stack of *n*_f_ = 80,000 frames, with *D*_av_ determined for respective *t_boxcar,n_* = 70, 105, 140, 175, 210 sec within that entire 280 s time range.

Using *n*_boxcar_ = 40,000 with Δ*n*_boxcar_ = 10,000 for successive overlapping boxcars, we performed ImFCS analysis on the same image stack of 80,000 frames as for Fig. 2A. The results are shown Fig. 2E-H (analogous to Fig. 2A-D). We find that CDFs of the *D* values obtained from these overlapping boxcars, from each complete data set (Fig. 2G) and from bootstrapped sampling of each (Fig. 2H), fall on top of one another confirming their indistinguishability. Likewise, the *D_av_* values determined from the sequential boxcars remain the same (0.7 μm^2^/sec), consistent with a resting steady-state (Fig. 2F). Left and right arrows in Fig. 2F point to the same Px units as those in Fig. 2B and reveal further fluctuations that were not resolved with non-overlapping boxcars. Despite small fluctuations, some Px units (e.g., indicated by left arrow) continue to exhibit relatively slow probe diffusion for a prolonged period, consistent with presence of diffusion-hindering membrane features (Fig. 2F). This evaluation shows that overlapping boxcar ImFCS analysis allows us to measure a *D*_av_ value from single cells every 35 sec. Further, consistent *D*_av_ values for PM-EGFP (Fig. 2F), as well as for EGFP-GG, YFP-GL-GT46, YFP-GL-GPI (Fig. S2), and Lyn-EGFP (data not shown), confirm that the plasma membrane of resting RBL cells is in a steady-state.

It is possible to decrease Δ*n*_boxcar_ thereby decreasing Δ*t*_boxcar_ and further reduce the time interval between two successive evaluation of *D*_av_ values. For example, Δ*n*_boxcar_ of 2,000 corresponds to Δ*t*_boxcar_ of 7 sec, with the trade-off that many more boxcar ImFCS analyses must be made to cover the time course of change. Such small Δ*t*_boxcar_ can be chosen when *D*_av_ of probe changes in tens-of-seconds time scale during a process. However, it is unnecessary to measure *D*_av_ every 7 sec if process occurs in the range of several minutes as for plasma membrane reorganization after Ag-stimulation (51, 52), as we further demonstrate in this study.

### Overlapping Boxcar ImFCS monitors stimulated diffusion changes in relevant timeframe

We evaluated the time of plasma membrane transition from resting (-Ag) to stimulated (+Ag) steady-states by measuring *D*_t_ (*D*_av_ as a function of time (*t_stim_*)) for PM-EGFP, EGFP-GG, and Lyn-EGFP before and after addition of Ag. We denote the averaged *D*_t_ in the resting (-Ag) condition as *D*_0_. After Ag-crosslinking of IgE-FcεRI, the ratio of *D*_av_ to *D*_0_ as function of time (*D*_t_/*D*_0_) represents the time-dependent change of *D*_av_ after stimulation. The ratio is determined for individual cells, thereby normalizing cell-to-cell variation of *D*_0_ (15). Based on our previous work (22), *D_t_*/*D*_0_ is expected to decrease with time for PM-EGFP and Lyn-EGFP and increase with time for EGFP-GG (Fig. 1B, Table 2). The shape of *D_t_*/*D*_0_ *vs t_stim_* plot reveals the dynamic characteristics of Ag-stimulated membrane reorganization sensed by a particular probe. We demonstrate the principle using PM-EGFP expressed in IgE-sensitized RBL cells.

**Table 2:** Membrane probe diffusion properties before, after, and during Ag stimulation as quantified with Boxcar ImFCS.

| Probe | Bag et al.<br>(22) | Present study using logistic model<sup>b</sup> |  |  |  |  |
| --- | --- | --- | --- | --- | --- | --- |
| | $\Delta[D_t/D_0]$<br>(%)<sup>a</sup> | $t_{on}$ [sec]<sup>d</sup> | $\Delta[D_t/D_0]$ (%)<sup>c</sup> | $t_{1/2}$ [sec]<sup>e</sup> | $k \cdot 10^3$<br>[sec<sup>-1</sup>]<sup>f</sup> | # data<br>points<br>used<sup>b</sup> |
| PM-EGFP | -8.0 |  | - 8.6 ± 1.7 | 735 ± 78 | 14 ± 9 | 19 out of 27 |
|  |  | 475 | - 8 | 719 | 9 | 27 (all<br>points) |
| EGFP-GG | 10.0 |  | 10.6 ± 0.5 | 562 ± 14 | 18 ± 3 | 19 out of 27 |
|  |  | 428 | 10.6 | 557 | 17 | 27<br>(all points) |
| Lyn-EGFP | -10.0 |  | - 9.4 ± 0.5 | 413 ± 14 | 20 ± 4 | 22 out of 33 |
|  |  | 298 | - 9.5 | 414 | 19 | 33 (all<br>points) |
| YFP-Syk | | | Stimulated<br>density,<br>($\rho_{Syk}^{Max}$) [μm<sup>-2</sup>] | | | |
|  |  |  | 0.020 ± 0.001 | 300 ± 21 | 9 ± 2 | 10 out of 16 |
|  |  | 53 | 0.020 | 298 | 9 | 16<br>(all points) |

<sup>a</sup> Steady-state values for $\Delta[D_t/D_0]$ determined previously with ImFCS from pooled cells in resting and stimulated states (22).
<sup>b</sup> Values of characteristic parameters determined from Boxcar ImFCS logistic transition model (Eqs. 1 and 2) and data plots shown in Fig. 4. Unshaded values are based on fitting all data points. Shaded values are based on bootstrapping with 30 trials on randomly sampled fraction of all data points as specified. ± values are standard deviation for bootstrapped parameters.
<sup>c</sup> $\Delta[D_t/D_0]$ : Percentage change of $D_{av}$ between stimulated steady state and resting steady state.
<sup>d</sup> $t_{on}$ : Onset time when the changes of $D_t/D_0$ becomes detectable by Boxcar ImFCS. This is defined as the time when the value of $D_t/D_0$ changes by 10% of the total difference between stimulated and resting steady-states, as determined by fitting all data points for each probe with Eq. 1.
<sup>e</sup> $t_{1/2}$ : Half-time for transition.

Ideally, individual single cells should be monitored for the entire time starting from the resting steady-state until an Ag-stimulated steady-state is reached such that *D*_av_ as a function of *t_stim_* covers both -/+ Ag steady-states and changes occurring during the transition. A typical single cell measurement following the entire process would take at least 20 min (280 sec for resting steady-state followed by at least 900 sec to reach Ag-stimulated steady-state), which is equivalent to recording about 340,000 frames. There are two major practical issues for such measurements: First, fluorescent probes inevitably undergo severe photobleaching due to continuous exposure to excitation beam for 20 min. Second, the configuration of the data acquisition system attached to instrumental setup must have the capacity 340,000 frames in one recording, and ours did not.

We addressed these practical limitations by evaluating seven sets of *D*_t_/*D*_0_ as a function of *t_stim_*on single cells in paired sets of resting and Ag-stimulated conditions, as shown in the general scheme of Fig. 3A. For each set, we record two 280 sec (*n_f_*= 80,000) image stacks of an ROI of a cell, before and during a selected time interval after Ag addition, performing Boxcar ImFCS in overlapping mode (*n*_boxcar_ = 40,000 and Δ*n*_boxcar_ = 10,000) for each interval. In this process we get five *D*_av_ values for each cell in the resting state (e.g., Fig. 2E,F), the average of which is defined as *D*_0_ for that cell. We set the time of Ag addition to be 0 sec. For each stimulated time interval, we collect an image stack corresponding to 280 sec, initiated at a specified time after Ag addition. In practice, we could make measurements only 60 sec after Ag addition, the delay due to time required for manual refocusing with our microscope set-up. The first set of stimulated measurements was done in the time interval of 60 to 340 (=60+280) sec after Ag addition. The *D*_t_ for this first time interval were determined at increasing *t*_boxcar_ (midpoint) corresponding to 130 (=60+70), 165 (=130+35), 200, 235, 270 sec after Ag addition. These time points represent the actual stimulation times (*t*_boxcar_ = *t*_stim_) from the time of Ag addition (*t*_stim_ is negative for measurements made in the resting state for each cell). For the first and all subsequent time intervals, *D*_t_ values at specified time points for each cell measured are divided by *D*_0_ for that cell to determine the normalized *D*_t_/*D*_0_ vs *t*_stim_ time course for that cell. For all time intervals *D*_t_/*D*_0_ vs *t*_stim_ measurements were repeated for 10-15 cells to achieve good statistics.

**Figure 3:**
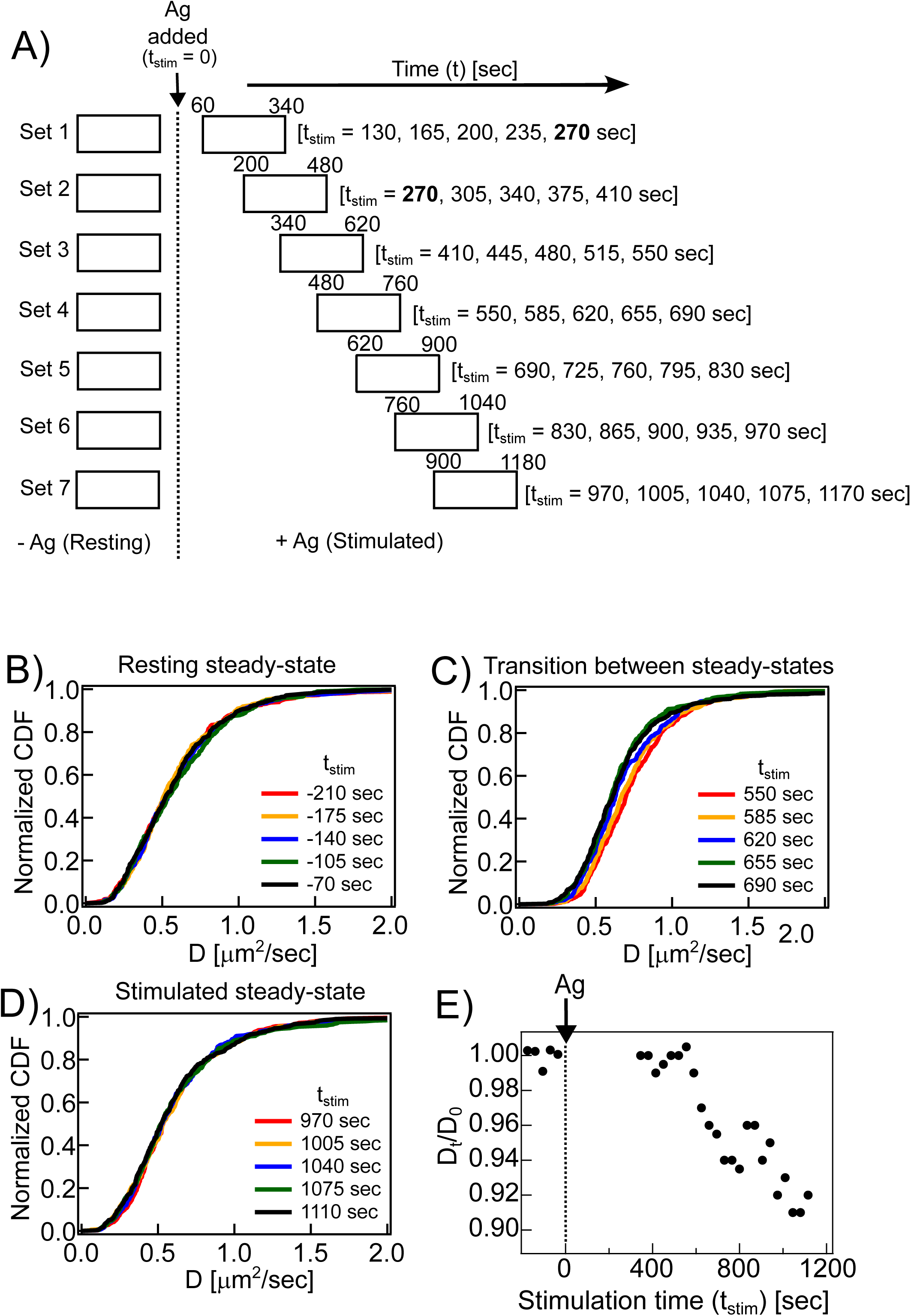
Overlapping boxcars on single cells at sequential time intervals quantify time-dependent change in probe diffusion. A) Boxcar ImFCS: Schematic of measurements to generate *D_t_/D_0_* against *t_stim_* plot for membrane probes. For each data set, one cell is measured for 280 sec before Ag addition to determine average diffusion coefficient at the resting steady state (*D_0_*), and the same cell is then evaluated for 280 sec at a set time interval after Ag addition to determine the average diffusion coefficient (*D_t_*) at corresponding *t_stim_*. For data set 1, +Ag measurements are carried out during 60-340 sec after Ag addition; the ImFCS analyses of overlapping boxcars (*n_boxcar_*=40,000, Δ*n*_boxcar_=10,000) yields *D_t_* at *t_stim_* boxcar midpoints of 130, 165, 200, 235, 270 sec. For data set 2, +Ag measurements are carried out during 200-480 sec after Ag addition, and analyses of overlapping boxcars yields *D_t_*at *t_stim_* of 270, 305, 340, 375, 410 sec. Data sets 3 - 7 are collected in the same manner on single cells, before Ag addition and at specified time intervals after Ag addition. B-D) Boxcar ImFCS analyses of cells expressing PM-EGFP after stimulation by Ag. CDFs of *D* values at specified time intervals related to time of Ag addition: B) resting steady-state, -280 – 0 sec; C) intermediate, transitional period after Ag addition (+480 – +760 sec); D) stimulated steady-state (+900 – +1040 sec). E) *D_t_/D_0_* against *t_stim_* plot for PM-EGFP probe obtained from measurements of many individual cells, following the scheme of part (A). Each data point represents the average *D_t_/D_0_*of 15-30 individual cells at specified *t_stim_*.

For the second time interval, we collected ImFCS data during 200 - 480 sec after Ag addition. Overlapping boxcar ImFCS analysis on this image stack yield *D*_t_ values at *t*_stim_ (midpoint) = 270, 305, 340, 375, 410 sec. As for all time intervals, measurements were also made on each cell in the resting state to determine that cell’s normalizing *D*_0_. *D*_t_/*D*_0_ was then determined as a function of *t*_stim_ for this second time interval. We continued to measure *D*_t_/*D*_0_ as a function of *t*_stim_ in for increasing time intervals after Ag addition. The complete set of seven time intervals correspond to: *t*_stim_ = 130-270 sec, 270-410 sec, 410-550 sec, 550-690 sec, 690-830 sec, 830-970 sec, and 970-1110 sec (Fig. 3A). The stimulated steady-state state was identified by unchanged *D*_t_/*D*_0_ during that time interval. This process was repeated for each probe (PM-EGFP, Lyn-EGFP, and EGFP-GG) to yield entire *D*_t_/*D*_0_ *vs t*_stim_ plots, which covered both resting and stimulated steady-states and the transition period between these steady-states. As noted, we measured 10-15 cells for each time interval for each probe, such that the entire *D*t/*D*0 versus *t*stim plot for each probe is based on data from 70-100 cells.

We exemplify our analysis scheme for PM-EGFP expressed in RBL cells, as shown in Fig. 3B-E. First, we checked the differences between resting (-Ag; t_stim_ < 0) and +Ag steady-states (*t*_stim_ = 910-1050 sec) **(Fig. 3A**). We previously observed ∼8% decrease of PM-EGFP’s *D*_av_, between -Ag and +Ag steady-states as determined from measurements on different pools of cells (22) (Fig. 1B). In the present study, we monitored the same individual cells in both steady-states and the results are strikingly similar. Time-dependent *D* CDFs for a representative cell in the resting state and 15 min after Ag addition (Figs. 3B and D, respectively) overlay each other, as expected for a membrane steady-states as sensed by PM-EGFP. Averaging together all the individual cells measured for these two conditions, the *D*_av_/*D*_0_ value decreases by ∼10% after Ag-stimulation (Fig. 3E).

We then evaluated the diffusion behavior of PM-EGFP in the intermediate time points after Ag stimulation, between the two steady-states. In initial measurements we found no significant change of *D*_t_ value (i.e., same as for the resting condition) in the first 360 sec after Ag addition, and we monitored *D*_t_/*D*_0_ at later times. As shown in Fig. 3E, we observed a decrease of *D*_t_/*D*_0_ value starting about 500 sec after Ag addition. The *D*_t_/*D*_0_ value continues to decrease over time until a plateau corresponding to the stimulated steady state is reached by about 1000 sec. The transition in *D*_t_ values can be seen for an individual cell by superimposing the *D* CDFs at increasing *t_stim_*between 630-770 sec (Fig. 3C). Unlike the group of *D* CDFs in the two steady-state conditions (-Ag/+Ag), these CDFs are distinguishable and shift to the left (decreasing *D*_t_) during the transition period. Thus, the transition between resting and stimulated steady-state for PM-EGFP diffusion occurs in the *t_stim_* = 500-1000 sec time frame. Overall, our results show three stages of stimulated membrane reorganization as sensed by PM-EGFP after Ag addition: i) a lag phase (0-∼500 sec) showing no detectable change in PM-EGFP’s diffusion as averaged over the membrane, ii) a transition phase (500-1000 sec) when membrane is detectably transitioning in organization such that the PM-EGFP’s averaged diffusion monotonically decreases, and iii) a stimulated steady state (>∼1000 sec) where PM-EGFP exhibits clearly slower diffusion.

Previous super-resolution microscopy of this inner leaflet Lo-preferring probe clearly demonstrated that nanoscale membrane ordering in proximity to Ag-clustered FcεRI is detectable within 5 min (20). Our ImFCS measurements extend this information by showing further that membrane reorganization continues globally on a longer timescale as Ag-crosslinking continues and the IgE-FcεRI clusters grow.

#### Fitting of time-dependent changes in probe diffusion with logistic model yields quantitative parameters that characterize transition between membrane steady-states

The observed shape of *D*_t_/*D*_0_ versus *t_stim_* for PM-EGFP is reminiscent of other processes such as dynamic changes in a population after a perturbation, which starts from an initial steady-state and undergoes a transition before establishing a final steady-state (53). Similar sigmoidal transitions between diffusive states were previously described in theoretical models of active matter that switches among various modes (54, 55). The kinetic data for these kinds of processes can be fitted empirically with a logistic model in the form of Eq. 1 to determine key parameters, such as the net change in the biophysical read-out between the initial and final steady-states (Δ[*D*_t_/*D*_0_]), the half-time for the transition (*t*_1/2_), and an apparent rate coefficient for the transition (*k*) indicating the steepness of the transition. The normalized value of *D*_t_/*D*_0_ at *t_stim_* = 0 is given by *C* whose fitted value should be about 1.

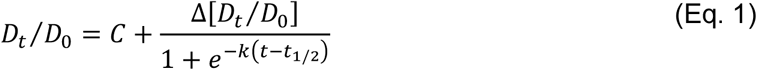

Fig. S4A illustrates the effect of each parameter on the shape of the transition curve for a probe that slows in diffusion after stimulation. By fitting (𝐷_𝑡_⁄𝐷_0_) data with this empirical model, we can quantitatively determine characteristic parameters for changes in a particular probe diffusion as driven by Ag-crosslinking of IgE-FcεRI.

Diffusion data for the example of PM-EGFP shown in Fig. 3E are reproduced in Fig. 4A, together with fits by the logistic model (Eq 1). The solid curve fits all 27 data points for *D*_t_/*D*_0_ *vs t_stim_*, each point averaged over 15-30 cells according to the scheme in Fig. 3A. The fitted values are: Δ[*D*_t_/*D*_0_] = -0.08 (i.e., 8% decrease), *t*_1/2_ = 719 sec, and *k* = 0.009 sec^-1^ (Table 2, shaded values). Because all the experimental *D*t/*D*0 vs *tstim* data are fit with a single curve, the fit parameters do not have error values. For error assessment we utilized bootstrapping: 30 randomly created subsets of the data points (each about two-thirds of all data points) were fitted individually with Eq 1, yielding the 30 gray lines shown in Fig 4A. The mean and standard deviation values of the fitted parameters from bootstrapped curves are provided in Table 2 (unshaded values). The values derived from the single fit of all 27 data points are the same as bootstrapped average within the standard deviation. The fitted value of Δ[*D*_t_/*D*_0_] for PM-EGFP corresponds to 8.6 ± 1.7% decrease in *D_av_* which is consistent with our previous report for this probe, which examined only the steady-states using measurements from different pools of cells (22) (Table 2). The characteristic half-time, *t*_1/2_ = 735 ± 78 sec (about 12 min), represents the time scale for global changes in membrane lipid organization caused by Ag-clustering of FcεRI as sensed by this probe. Notably, the low *k* value (0.014 ± 0.009 sec^-1^) quantifies our observation that the transition of PM-EGFP diffusion from resting to stimulated steady-state is slow and gradual.

**Figure 4:**
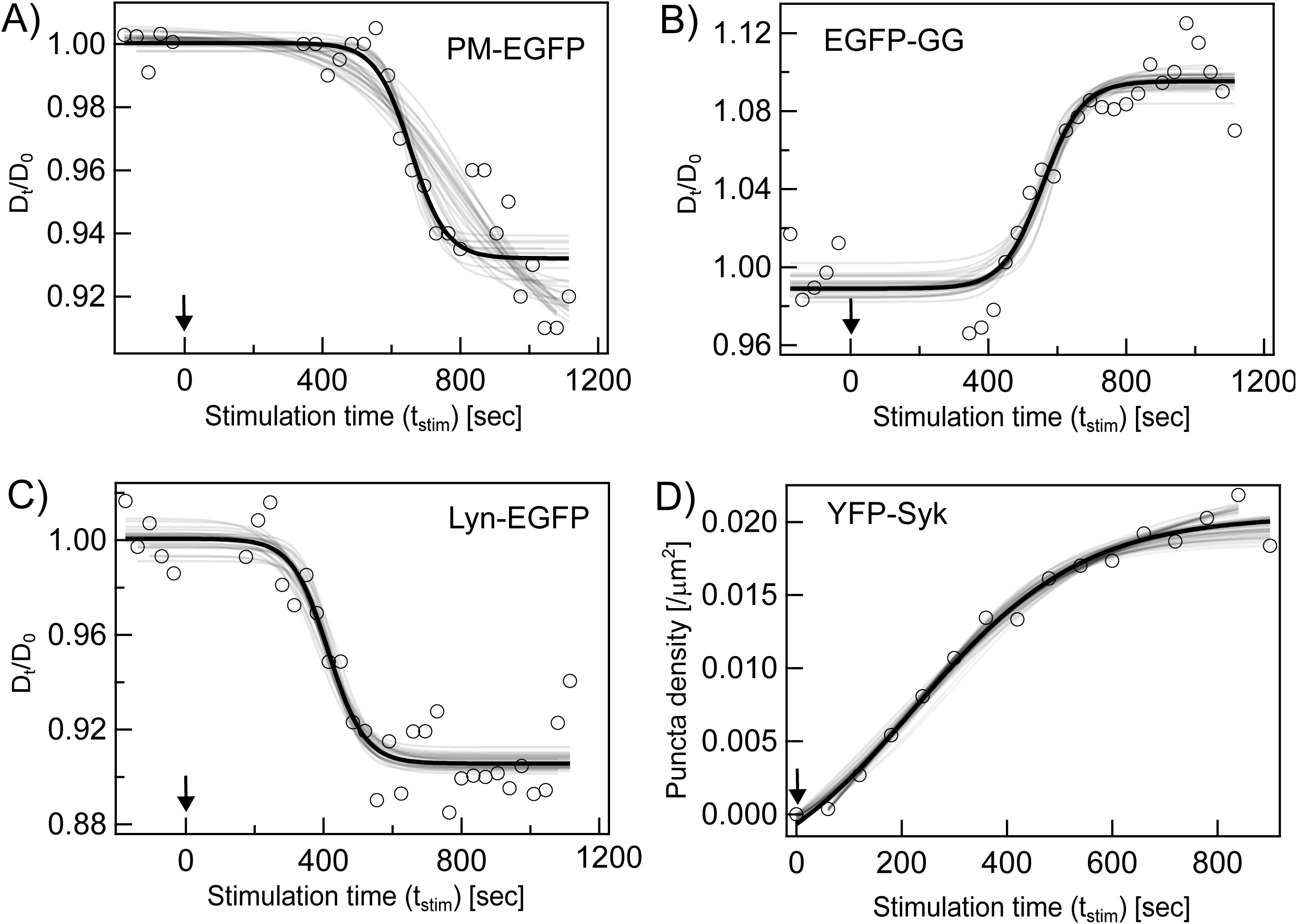
Probe diffusion and signaling data are well-fit by logistic model. A – C) Boxcar ImFCS analyses of specified membrane probe: A) PM-EGFP; B) EGFP-GG; C) Lyn-EGFP. Each data point represents the average *D_t_/D_0_* of 15-30 individual cells at specified *t_stim_*. D) Time course for appearance of Syk puncta; each point represents the average of 12 cells (29). Arrow indicates the time of Ag addition in each case. Data are fit with logistic model (Eq. 1 for A-C, Eq. 2 for D) using two approaches: Black lines are fits when all data points are included, and gray lines are fits to bootstrapped subsets of data points. Parameters derived from fits are provided in Table 2.

Because we chose the logistic model heuristically, we evaluated the fitted parameters by simulation (Fig. S5 and Table S1). We simulated image stacks (128 Px units, 700 sec) using particles with parameters for changes in diffusion coefficients set by Eq. 1. We selected values for Δ[*D*_t_/*D*_0_], *t*_1/2_, and *k* like those obtained for PM-EGFP (Table 2) and found that fitted values of all parameters thus obtained match the simulated values (Table S1). Further simulations confirmed that Δ[*D*_t_/*D*_0_] and *t*_1/2_ are accurately recovered under all tested conditions. We found the *k* value to be accurately estimated for simulated *k* ≤ 0.05 sec^-1^, which is the range we observed experimentally for PM-EGFP, EGFP-GG, and Lyn-EGFP, as described in the next section.

### Part II: Boxcar ImFCS to Evaluate Dynamic Changes in Membrane Organization with Differentially Selective Probes

#### The kinetics of stimulated diffusion changes for selective membrane probes are distinctive

After establishing the overlapping Boxcar ImFCS analysis using the Lo-preferring lipid probe, PM-EGFP (Part I), we examined stimulated changes in Lyn-EGFP and EGFP-GG. The tyrosine kinase, Lyn, involved in the first steps of FcεRI-mediated signaling, is a Lo-preferring, proteolipid with the same palmitate/myristoylate membrane anchor as PM-EGFP (Fig. 1A). Lyn also has substantial protein composition including SH2, SH3, and kinase domains. EGFP-GG is an Ld-preferring lipid probe, based on the K-RAS membrane anchor and geranylgeranylation (26) (Fig. 1A). The experimental and analysis protocols for these two probes were carried out as for PM-EGFP (Fig. 3A). The resulting plots of *D*_t_/*D*_0_ *vs t_stim_*for all three probes are compared in Fig. 4A-C. For each probe the entire data set (solid curve) and bootstrapped random subsets (gray curves) are fit with the logistic model (Eq. 1). The parameters derived from all of these fits are provided in Table 2, with shaded values corresponding to solid curve fits and unshaded values corresponding to bootstrapped curve fits. The values determined for each probe from both fits are consistent within the standard deviations provided by the bootstrapped fits. Table 2 also includes our previous steady-state measurements with each of these probes using pooled samples (22). The results are consistent: Δ[*D*_t_/*D*_0_] is about -0.1 for both Lyn-EGFP and PM-EGFP and about +0.1 for EGFP-GG, corresponding to about 10% decrease or increase of respective *D_av_* values in the stimulated steady-state after addition of Ag to cluster FcεRI.

Boxcar ImFCS measurements of all three probes, as averaged over multiple single cells, revealed kinetic details of the transition between resting and stimulated steady-states with parameters *k* and *t*_1/2_. The low *k* values for all three probes (0.01 – 0.02 sec^-1^) reflect slow and gradual transitions from resting to stimulated steady-states for all PM-EGFP, Lyn-EGFP, and EGFP-GG (Fig. 4A-C). Interestingly, we found significant different values for *t*_1/2_, and these represent the time scale of reorganization of the membrane environment as selectively sensed by each probe. Lyn-EGFP with selective lipid and protein components responds more quickly (*t*_1/2_ = 413 ± 14 sec) than the other probes after Ag stimulation. Comparing the lipid probes, disorder-preferring EGFP-GG senses membrane reorganization faster (*t*_1/2_ = 562 ± 14 sec) than order-preferring PM-EGFP (*t*_1/2_ = 735 ± 78 sec). The contrasting times for diffusional transition after Ag stimulation for the three probes (*t*_1/2_ ≃ 410 – 750 sec) reflects their respective lipid-based and protein-based modes of membrane interactions. We also used the logistic model to determine a value for *t*_on_, corresponding to a the first detectable change in *D*_t_/*D*_0_ (10% of Δ[*D*_t_/*D*_0_]), for each probe (Table 2). These parallel the respective *t*_1/2_ values. Thus, following a differential lag period, all three probes undergo a small (± 10%) and gradual (*k* = 0.01-0.02 sec^-1^) diffusional shift after Ag stimulation, with Lyn-EGFP significantly preceding the two lipid probes.

#### Stimulated Syk recruitment is detectably faster than global changes in membrane probe diffusion

To explore further the cell biological context of our measurements of membrane probe diffusion we compared the time scale (*t*_1/2_) of a stimulated functional response. The signaling step just after phosphorylation of Ag-crosslinked FcεRI by Lyn kinase is recruitment of cytoplasmic Syk kinase to the plasma membrane where it phosphorylates its protein substrates (28). We previously used TIRFM imaging to quantify Syk puncta appearing on the plasma membrane as a function of time after antigen stimulation (29). We used these data points to evaluate the kinetics of Syk recruitment in a similar manner as described for the stimulated time-dependent changes of probe diffusion. We applied a logistic model, Eq. 2, to fit the time dependent changes of Syk puncta density (𝜌_*Syk*_(𝑡)) at the plasma membrane,

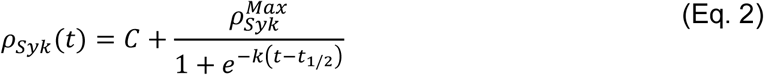

where *ρ*_*Syk*_^*max*^ is the saturated value of YFP-Syk puncta density, *t*_1/2_ is the half-time for the transition, and *k* is an apparent rate coefficient indicating the steepness of the transition. The value of *ρ*_*Syk*_ at *t_stim_* = 0 is given by *C* whose fitted value should be about 0.

As shown in Fig. 4D and Table 2, Syk puncta appear within a minute after Ag addition (*t*_on_ = 53 sec, Table 2), and the puncta density (*ρ*_*Syk*_) increases non-linearly with time until a steady-state of ∼ 0.020 μm^-2^ is reached (data points from (29) in Fig. 4D). This curve clearly contrasts with the stimulated diffusion changes of the tested membrane probes for which there is a ≥ 5 min delay before any change is detected by ImFCS (Fig, 4A-C; *t*_on_, Table 2). Syk recruitment kinetics has a *t*_1/2_ = 300 ± 21 sec and *k* = 0.009 ± 0.002 sec^-1^. The results quantify stimulated Syk recruitment occurring as an gradual process after a relatively short lag period. Thus, stimulated Syk recruitment and global membrane reorganization (as most directly probed by diffusion of PM-EGFP and EGFP-GG) occur with different time dependence. Overall results are consistent with the view that Ag-crosslinking of IgE-FcεRI causes nanoscopic ordering of the membrane lipid environment proximal to the minimally clustered FcεRI to initiate transmembrane signaling and recruitment of cytosolic Syk within minutes (20, 29). As Ag-crosslinking continues to increase the size of FcεRI clusters, the membrane segregation into Lo-like and Ld-like regions continues until it reaches a new steady-state (Fig. 4).

## DISCUSSION

The functional impact of Lo-like nanodomains in the plasma membrane, also called rafts, have been established in a variety of biological processes using high resolution fluorescence microscopy approaches. The processes investigated include transmembrane signaling mediated by IgE receptor FcεRI (20, 22), B cell receptor (21), T cell receptor (56–58) and receptor tyrosine kinases (59) as well as endocytosis (60), and viral budding (61). ImFCS has proven to be a valuable and versatile method for evaluating membrane organization by measuring diffusion coefficients of selective membrane probes. Because of the exceptional precision possible from very large data sets, measurements can detect subtle changes in probe diffusion due to modulation of the membrane environment caused by a perturbation, such as Ag stimulation of immune cells. In general, ImFCS has been used to examine stationary systems (steady-states). As described herein we devised overlapping Boxcar ImFCS, based on raw data segmentation, to evaluate membrane organization in a steady-state as well as changes occurring as the system moves from one steady-state (resting, -Ag) to another (stimulated, +Ag). By fitting the time-dependent change with a logistic model (Eq. 1) we extract kinetic parameters that quantitatively characterize the membrane reorganization as experienced by selective probes.

### Boxcar ImFCS with bootstrapping evaluates a steady-state

The steady-state of a system is defined by unchanged dynamic properties as averaged over time. As previously introduced (15), non-overlapping Boxcar ImFCS can demonstrate the resting steady-state of plasma membrane (Fig. 2). The values for *D_a_*_v_ should be the same in the steady-state condition, but paired students’ t-test between the *D* distributions may still indicate a significant difference if a very large number of data points, such as with our measurements (62). Likewise, the CDFs of *D* values from large boxcars measuring the same steady-state may not overlay completely.

More confidence can be gained by considering overlapping boxcars generated from the same image stack. We see that CDFs of *D* values obtained from five overlapping boxcars do overlay (Fig. 2) and pairwise t-test consistently yields non-significant p-values. Bootstrapping of random sub-samples provides additional confidence for both non-overlapping and overlapping boxcars by qualitatively indicating whether the *D* CDFs of sub-samples overlay each other (Fig. 2). Our evaluation of resting RBL cells in this manner for multiple probes with different types of membrane interactions demonstrated a resting steady-state (Fig. S2).

### Stimulated membrane reorganization is sensed differentially by distinctively selective probes

Previous studies established that Ag (DNP-BSA) crosslinking of anti-DNP IgE-FcεRI in RBL cells stabilizes proximally an ordered lipid environment in the inner leaflet, into which order-preferring Lyn kinase partitions preferentially to initiate transmembrane signaling (20, 26, 52, 63, 64). Consequent phosphorylation of FcεRI by Lyn recruits cytoplasmic Syk kinase, followed by assembly of signaling proteins, and ultimately degranulation as a cellular response to the antigenic stimulus (28). Ag-induced FcεRI co-clustering together with Lyn and order-preferring lipids (e.g., PM-EGFP) in the first few minutes has been measured on the nanoscale with super resolution fluorescence localization microscopy (51). ImFCS, which measures diffusion coefficients (*D)* of probes in individual Px units (size: 0.1 μm^2^) to provide a *D* map which can be averaged across a 40 - 60 μm^2^ region of the membrane (*D_av_*) (Fig. S1). We previously showed with pooled cells that that the *D_av_* of order-preferring Lyn-EGFP and PM-EGFP and disorder-preferring EGFP-GG are different before addition of Ag to crosslink IgE-FcεRI and after the final steady-state is reached at 15 min (Fig. 1; (22)). In the present study, Boxcar ImFCS on individual cells before, during, and after Ag-stimulation delineates the transition between resting and stimulated steady-states. This time-dependent process occurs more slowly than the initial nanoscale rearrangement of crosslinked IgE-FcεRI (51) and reflects global membrane reorganization and raft condensation around larger nanoclusters of crosslinked receptors.

Xu et al. (65) previously determined that the crosslinking capacity of commonly used Ag, DNP-BSA (averaged stoichiometry ∼15 DNP/BSA) is not simply defined because presentation for binding to anti-DNP IgE of the hydrophobic moiety DNP is restricted by its level of exposure on the BSA surface. The time-dependent binding to IgE is well-fit by a “slow hapten exposure” model, yielding forward/reverse binding rate constants and other parameters including an equilibrium constant for hapten exposure. Notably, about one (on average) DNP is initially exposed on the protein surface and this DNP hapten binds relatively quickly. This monovalent binding is followed by slower crosslinking of IgE sites by slowly exposed DNP haptens.

Comparing the DNP-BSA binding curve to phosphorylation of FcεRI by Lyn kinase, the time-dependent binding and phosphorylation curves were found to overlap at all but the earliest time points and only as crosslinking of IgE-FcεRI into clusters proceeds.

Using conditions like those in the present study, Shelby et al. (51) used super resolution microscopy to quantify time-dependent aggregation of IgE-FcεRI after addition of DNP-BSA. They monitored both cluster size using auto-correlation functions and diffusion coefficients using single particle tracking (Figs. 1-3 in (51)). Plotting this diffusion coefficient *vs* number of IgE-FcεRI per cluster (*N*) with increasing time, revealed two distinct stages of receptor clustering. In the first stage, which precedes stimulated Ca^2+^ mobilization (0 – ∼1 min after Ag addition), IgE-FcεRI diffusion slows markedly as *N* increases to ∼20. In the second stage (2 - 15 min), IgE-FcεRI diffusion decreases much more gradually as *N* increases from ∼40 to ∼80. A steady-state, observed as distinct puncta, occurs by 15 min. Considering also the previous DNP-BSA binding studies (65), led to the conclusion that the initial decrease in IgE-FcεRI diffusion and the signaling responses observed at early stimulation times occur concurrently with the formation of small clusters, which continue to grow in size as they become more heavily cross-linked. Our Boxcar ImFCS measurements of *D_av_* become sensitive to diffusion changes of non-receptor membrane probes (Fig. 1) at the later stage and monitor membrane reorganization until the new steady-state of clustered IgE-FcεRI puncta is reached.

The probes we evaluated, Lyn-EGFP, PM-EGFP, and EGFP-GG are all lipid-anchored to the plasma membrane inner leaflet and have differentially selective membrane interactions (Fig. 1). They exhibit distinctive diffusional behaviors as revealed by the Boxcar ImFCS measurements (Fig. 4). The respective *D_t_/D_0_ vs t_stim_* curves are quantified by a logistic model (Eq. 1) to yield values for the net change in diffusion coefficient Δ[*D_t_/D_0_*], the intrinsic transition rate coefficient *k* (steepness of the transition curve), and the half-time for the transition (*t_1/2_*). The lag time before the detected diffusion change can be quantified as *t_on_*(10% of Δ[D_t_/D_0_]). Fitting all time course data yields a single averaged value for each of these parameters, and bootstrapping 30 random sub-samples of data points yields values averaged over sub-fits together with an error estimate (Table 2). The Δ[*D_t_/D_0_*] value reflects the net change in membrane environment sensed by a probe as it diffuses in regions that are stabilized in the steady-state after Ag addition. Values are negative for Lyn-EGFP (-9.4 ± 0.5) and PM-EGFP (-8.6 ± 1.7) indicating a more Lo-like environment in which these order-preferring probes partition and diffuse slower. The value is positive for EGFP-GG (+10.6 ± 0.5) indicating a more Ld-like environment in which this disorder-preferring probe partitions and diffuses faster. These results agree with previous ImFCS measurements averaged over pooled cells in just the two steady-states ((22); Table 2), and they support the view that Ag-crosslinking of IgE-FcεRI into large clusters causes stabilization of Lo-like rafts and phase-like separation of Ld-like regions within the membrane inner leaflet. Transmembrane signaling is initiated with small IgE-FcεRI clusters and lipid rearrangement on the nanoscale (20, 51), and this later manifestation of membrane organization may be an amplification and stabilization of the early stage as the crosslinked receptor clusters continue to grow in size. The later stage of membrane organization may also play a role in downstream signaling leading to degranulation and/or endocytosis, neither of which goes to completion under our experimental conditions room temperature.

Table 2 shows that the three membrane probes are distinguished by their t_on_ and t_1/2_ values, with Lyn-EGFP (298 sec and 414 sec, respectively) < EGFP-GG (428 sec and 557 sec) < PM-EGFP (475 sec and 719 sec). Drawing from the data reported in Shelby et al.(51) as described above, the t_on_ values correspond roughly to IgE-FcεRI cluster sizes (*N*) for Lyn of 49, for EGFP-GG of 56, and for PM-EGFP of 58. The *t_1/2_* values correspond roughly to cluster sizes (*N*) for Lyn of 55, for EGFP-GG of 63, and for PM-EGFP of 71. Because Lyn-EGFP and PM-EGFP have the same palmitate/myristoylate lipid anchors (Fig. 1) the striking difference between these two probes can be ascribed to Lyn’s additional protein domains, SH2, SH3, and kinase, which associate with cytoplasmic protein segments of FcεRI (66). We showed previously that its lipid anchor causes Lyn-EGFP to partition first into ordered-lipid regions surrounding clustered IgE-FcεRI followed by the protein-based interactions. In contrast, S15-Lyn, which is anchored to the membrane with a disorder-preferring lipid anchor, does not exhibit a decreased diffusion coefficient and does not phosphorylate FcεRI upon Ag addition (22). The smaller t_1/2_ and N values for Lyn-EGFP compared PM-EGFP indicate that partitioning of Lyn into ordered regions beginning within a minute (20) facilitates the slower diffusion caused by subsequent protein interactions that thereby becomes detectable earlier. The change in PM-EGFP diffusion is not detectable until the IgE-FcεRI are sufficiently large to stabilize a larger ordered (raft-like) environment.This trend is consistent with previous studies investigating sensitized RBL cells interacting with micropatterned Ag surfaces (52): As visualized with fluorescence microscopy, Lyn-EGFP was detectably recruited to the Ag-crosslinked IgE-FcεRI patches before PM-EGFP was detectably recruited.

The t_1/2_ value is significantly less EGFP-GG (∼9 min) than that for PM-EGFP (∼12 min). This difference suggests that the unsaturated GG-lipids begin to move faster in more disordered environments before the ordered regions are sufficiently stabilized to comparably slow the saturated PM-lipids. These values are both in the range of predictions from computer simulations, which estimate lipid reorganization and coalescence of membrane nanodomains into stable structures occurring in ∼10-15 min (67–69). As investigated for cells in a resting steady-state, the inner leaflet of the plasma membrane is generally disordered with a greater abundance of Ld-preferring lipids (15, 70). It is reasonable that the sparser distribution of Lo-preferring lipids takes longer to dynamically assemble around the clustered IgE-FcεRI after Ag addition. Among other factors, the cortical cytoskeleton may participate to stabilize Lo/Ld phase-like separation (14). For example, when sensitized cells were placed on micropatterned Ag surfaces, actin was observed to be recruited to IgE-FcεRI patches after Lyn-EGFP and prior to PM-EGFP (52).

Interestingly, once the diffusion transition begins for each of these three probes the *k* values are roughly the same (within error) for Lyn-EGFP (0.019 sec^-1^), PM-EGFP (0.009 sec^-1^), and EGFP-GG (0.017 sec^-1^) (Table 2). In all cases the transition is slow and gradual in contrast to an abrupt switch-like process expected for *k* values larger than 0.2 sec^-1^ (Fig. S5). Agreement with the logistic model indicates a cooperative diffusional change for all three membrane probes and the similarly gradual change may be related to dependence on the IgE-FcεRI cluster size.

To compare the diffusion transition of these three membrane probes to early signaling events, we evaluated data for Ag-stimulated recruitment of cytoplasmic Syk kinase to the plasma membrane (29). We found that these data are well fit with a logistic model (Eq. 2). In this case, no membrane-associated Syk is detectable in the absence of Ag, and the net change is fit as “stimulated puncta density (μm^-2^),” with values for *t_on_*, *t_1/2_*, and *k* extracted as for the membrane diffusion probes (Table 2). Stimulated Syk recruitment initiates with nanoscale clusters of IgE-FcεRI (71), yielding measurable *t_on_*= 53 sec and *t_1/2_* = 298 (corresponding to *N* = 20 and 49, respectively (51) for appearance of puncta. It is not surprising that these values are much shorter than those for diffusion change of the three membrane probes which become detectable when the IgE-FcεRI clusters are larger (Table 2). In essence, nanoscale stabilization of rafts around the smaller receptor nanoclusters (*N* ∼ 20) is sufficient to initiate transmembrane signaling while the physical properties of the plasma membrane continue to change as the nanoclusters grow larger (*N* ≳ 50) making phase-like organization progressively more stable leading to noticeable changes of probe diffusion at sub-micron length scale. The extracted *k* value for Syk recruitment is in the same range as for the three membrane probe diffusional changes, consistent with recruitment continuing as the receptor cluster size increases.

#### Simulations show that Boxcar ImFCS and Logistic Model yield robust parameter values for present experimental system but has limitations

We developed overlapping Boxcar ImFCS to measure changing diffusion of selected membrane probes after Ag addition to cluster IgE-FcεRI. We set conditions appropriate for biological and technical aspects of our experimental system, which included the rate at which process occurs (biological) and limitations of our microscope set-up (technical). Our choices of *n_boxcar_* = 40,000 frames was based on requiring sufficiently high quality ACFs (Fig. S3) and our choice of *Δn_boxcar_* = 10,000 frames (35 sec) was based on the intrinsic time scale of the process, as previously established (22). To evaluate capacity and limitations of these choices, we carried out simulations as described in the and represented in Fig. S5 and Table S1. We addressed the “drag effect” imposed by our selected size of *nboxcar* due to averaging over the entire 40,000 frames to yield a *D* value for an assigned *t_boxcar_* (or *t_stim_*) which is determined by both *n_boxcar_* and *Δn_boxcar_*(Figs. 2 and 3). This drag can effectively reduce the simulated transition rate coefficient *k* (steepness) compared to the theoretical curve dictated by the input values (Eq. 1). We also evaluated the effect of magnitude of the overall change (*Δ[D_t_/D_0_]*) to consider whether a change larger than the ∼10% we observe would make a significant difference. Carrying out simulations with input values like those determined for PM-EGFP, then changing values for *k*, we made several comparisons as shown in Fig. S5 and Table S1. The simulations show that our choices of *n_boxcar_*

= 40,000 and *Δn_boxcar_* = 10,000 are sufficient for determining robust values of *Δ[D_t_/D_0_]* and *t_1/2_* for both *Δ[D_t_/D_0_]* = 10% and 90%. These choices are also sufficient for *k* values in the range of 0.01 sec^-1^ and 0.05 sec^-1^, corresponding to slowly gradual transitions. However, they result in marked deviations for *k* = 0.20 sec^-1^, corresponding to a steep transition, due to the drag of averaging over the entire *n_boxcar_* (Fig. S5). To capture such steep (more switch-like) transitions would require correspondingly shorter *n_boxcar_*, which would also require brighter, more photostable fluorophores for higher quality ACFs (Fig. S3) and many more boxcar analyses to cover the time course. We can confidently conclude that our choices of *n_boxcar_* and *Δn_boxcar_* are adequate for our experimental system because the diffusional transitions of the three membrane probes responding to Ag-stimulated membrane reorganization are delayed (*t_1/2_* = 400-700 sec) and gradual (*k* = 0.01 – 0.05 sec^-2^ (Table 2).

#### Future refinements will enhance Boxcar ImFCS capacity to evaluate stimulated changes in membrane organization

Our development of Boxcar ImFCS within our experimental conditions gave us the novel opportunity to characterize membrane reorganization occurring on minutes time scale as mediated by clustered IgE-FcεRI after Ag stimulation.

However, these measurements had practical limitations as described in previous sections Ideally, a 20 min long measurement could be carried out on single cells to get *D_t_/D_0_* against *t_stim_* plots for each probe tested. This would be 5 min in resting conditions and continuous 15 min after Ag addition corresponding to recording of 343,000 frames (at 3.5 ms/frame speed). Our technical limitations, including photostability of fluorophores and limited computer memory, precluded such long measurements. We overcame these limitations with a labor-intensive alternative (Fig. 3) requiring measurements on a total of about 100 cells per probe (∼15 cells from three biological replicates for each set). With 20 min long, single cell measurements, the equivalent level of data statistics could be achieved by measuring a total of ∼15 cells.

Current limitations of Boxcar ImFCS can be addressed for a broad range of applications. For example, it was recently reported that *D* values of membrane probes can be determined with good precision in ImFCS analysis using only 2,500 frames together with machine learning (72). With this approach and data acquisition at 3.5 ms/frame it would be possible to use non-overlapping boxcars and measure *D_t_* every 8.75 sec = *Δt_boxcar_*. This would yield 7 data points per minute to construct *D_t_/D_0_ vs t_stim_* curve, sufficient for measuring diffusional changes on sub-minute time scale without using overlapping boxcars. The *Δt_boxcar_*can be decreased further by using overlapping boxcars to evaluate even faster diffusional changes. Overlapping boxcars may introduce the problem of “drag” on the apparent rate of transition, such as for evaluation of *k* with the logistic models as discussed above. Although use of machine learning is attractive, synthesizing the training data set for ImFCS data analysis generally requires understanding of the underlying modes of diffusion processes; for example, hindered diffusion caused by various forms of membrane heterogeneity features (73, 74).

To address the limitations of photostability, fluorescent dye technology is rapidly evolving to create probes for long-term live cell microscopic measurements. These could allow single cell measurements lasting as long as the 20 minutes required in our study of membrane reorganization. For example, monomeric variants of StayGold (75), shown to be promising bright alternative to EGFP, were recently used in multiplexed FCS measurements (76). Similarly, bright and photostable organic probes such as the series of Jenelia Fluor (JF) dyes (77) can be used for long-term measurements. As for any probe, care must be taken to ensure retained function of the proteins and lipids conjugated to these fluorophores.

Typically for fluorescence microscopy measurements, a sample must be refocused when a stimulus (e.g., Ag) is manually added to the dish mounted on the microscope stage, and this delays the first post-stimulation measurement. This may be improved if the stimulant is introduced through perfusion channels in a microfluidic device causing minimal disruption of the sample. As recently reported (78), it is possible to calculate ACFs in real time during ImFCS data acquisition. With this approach any distortions in ACFs during data acquisition, such as arise from sample defocusing, can be easily detected. The stimulant can be introduced as the acquisition proceeds until undistorted ACFs are observed for the earliest possible data point after stimulation. Although we were limited by data storage capacity, large amount of raw data (e.g, 343,000 frames) can now be recorded by modern GPU-based parallel computing or CPUs with 64 GB or more RAM. These enhanced capacities would also significantly accelerate data analysis speed.

## DATA AVAILABILITY

Raw data and simulation codes are available from the corresponding author upon reasonable request.

## ACKNOWLEDGEMENTS

We thank Alice Wagenknecht-Wiesner (Cornell University) for assisting with the preparation of constructs. This work is supported by the National Institute of General Medical Sciences (NIGMS) Grant R01GM117552. N.B. acknowledges SRG funding from Anusandhan National Research Foundation (ANRF) (SRG/2023/001423) for financial support during manuscript preparation.

## AUTHOR CONTRIBUTIONS

G-S.Y., N.B., and B.A.B. conceived and designed the research project; G-S.Y. performed all experiments and simulations; N.B. developed key technical concepts; G-S.Y., N.B., and B.A.B. analyzed data; and G-S.Y., N.B., and B.A.B. wrote the paper.

