## Supplemental information for "Boxcar Imaging FCS Reveals Membrane Raft Stabilization Kinetics in Antigen-Stimulated Mast Cells"

### Materials and Experimental Methods:

#### Reagents

Minimum essential medium (MEM), Opti-MEM, Trypsin-EDTA (0.01%), and gentamicin sulfate were purchased from Life Technologies (Carlsbad, CA). Fetal Bovine Serum (FBS) was purchased from Atlanta Biologicals (Atlanta, GA). The multivalent antigen (Ag), DNP-BSA, was prepared by conjugating DNP sulfonate (Sigma-Aldrich) to bovine serum albumin (BSA), yielding an averaged stoichiometry of ~15 DNP/BSA (1). Phorbol 12,13-dibutyrate (PDB) was obtained from Sigma-Aldrich (St. Louis, MO). Stock solution of PDB was prepared in DMSO and stored at -80°C.

#### Cell culture, chemical transfection, sensitization, and stimulation

RBL-2H3 cells were grown and maintained in T-25 flask in full growth medium containing 80% MEM, 20% FBS, and 10 mg/L gentamicin sulfate at 37°C and 5% (v/v) $CO_2$ environment. Cells were transfected with FuGENE HD transfection kit (Promega) using 0.5-1 µg plasmid per 35-mm dish. The details of transfection protocol are described elsewhere (2). The chemically transfected plasmids used in this study encode the following proteins: Lyn-EGFP (3), PM-EGFP(3) , EGFP-GG (3), YFP-Syk (4).

On the day of ImFCS measurements, the transfected cells in a 35-mm dish were washed with buffered salt solution (BSS: 135 mM NaCl, 5.0 mM KCl, 1.8 mM $CaCl_2$, 1.0 mM $MgCl_2$, 5.6 mM glucose, and 20 mM HEPES; pH 7.4) twice (1 mL each time) followed by addition of 1 mL of anti-DNP IgE (2 µg/mL) prepared in BSS for 45 min at room temperature. The 35-mm dish was then mounted on the TIRF microscope and an ImFCS measurement (see below) was done on a suitable cell to determine the $D_{av}$ value in resting state. This was followed by Ag stimulation by adding 0.5 µg/mL of DNP-BSA directly to this dish and then ImFCS measurements were done at the specified time (stimulation time, $t_{stim}$, in Figs. 3 and 4, Main Text).

**Total internal reflection fluorescence microscopy (TIRFM) and Imaging FCS (ImFCS) measurements:**

A home-built TIRFM system with a 100x/1.49NA oil immersion objective and an EMCCD camera attached on the side port was used for all ImFCS measurements (2). A 488 nm laser was used for excitation of the sample. A suitable transfected cell in a 35-mm MatTek dish was chosen and a small region of interest (ROI) of typically 25×25 pixel units (Px units, dimension 320 nm) on the cell was selected. A movie of 80,000 frames from this ROI was collected and further used for various modes of Boxcar ImFCS analyses as described in the Main Text. The autocorrelation functions (ACFs) obtained for each Px unit for a given boxcar were analyzed to generate a map of diffusion coefficient values (*D* map) for that boxcar using a FIJI/ImageJ plug-in (Imaging_FCS 1.491; available at http://www.dbs.nus.edu.sg/lab/BFL/imfcs_image_j_plugin.html) following the established protocols (5–7). All imaging measurements were done at room temperature.

**Supplemental Figures (Experimental):**

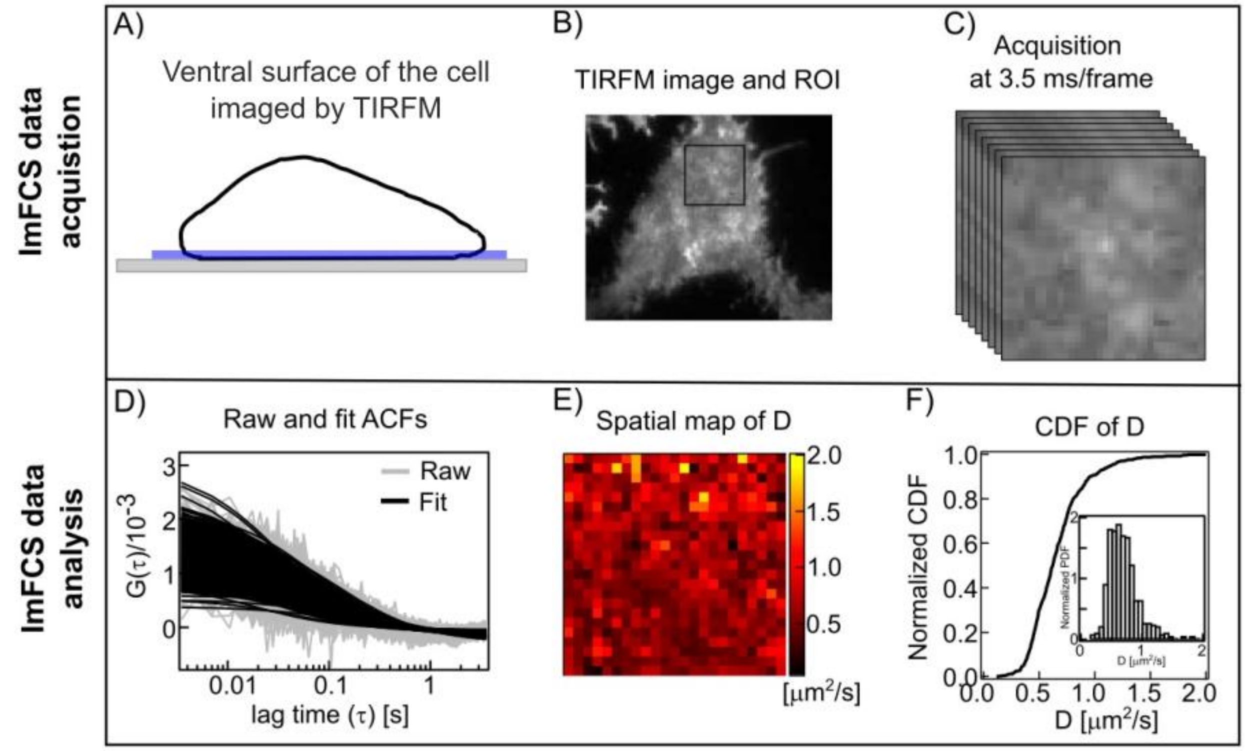


**Figure S1:** <u>Diffusion coefficients are measured in individual Px units and averaged over ROI.</u> Work flow of ImFCS data acquisition and analysis is illustrated with PM-EGFP expressed in RBL cell: A-B) A region of interest (ROI) including 20×20 Px units (or larger) is selected on the ventral surface of the fluorescently labeled cell. C) An image stack of 80,000 frames at a speed of 3.5 ms per frame is recorded. D) Shown are raw ACFs (grey) generated from fluorescence fluctuations at each Px unit and the corresponding fits (black) to determine values of respective diffusion coefficients ($D$, $\mu m^2/s$). E) The $D$ values of each Px unit obtained after fitting can be presented as spatial maps. F) The $D$ values can be pooled to generate a normalized cumulative distribution function (CDF) or probability distribution function (PDF, inset).

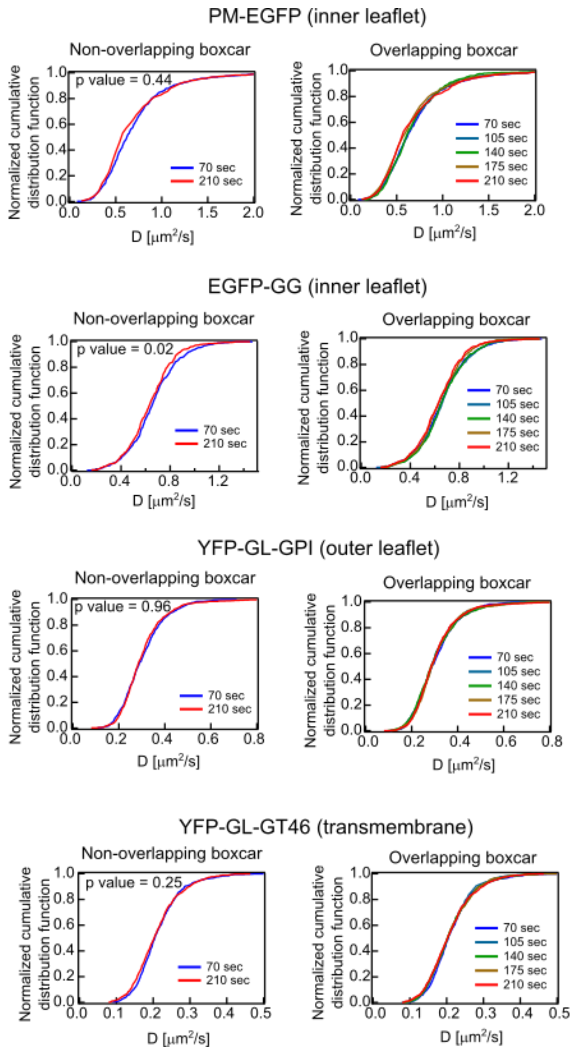


**Figure S2**: Normalized *D* CDFs for representative non-functional membrane probes in resting RBL cells. For both non-overlapping (left panels) and overlapping (right panels) boxcars, steady-state diffusion is confirmed by curves that overlay for each of YFP-GL-GT46 (transmembrane protein), YFP-GL-GPI (outer leaflet lipid probe), PM-EGFP and EGFP-GG (both inner leaflet lipid probes).

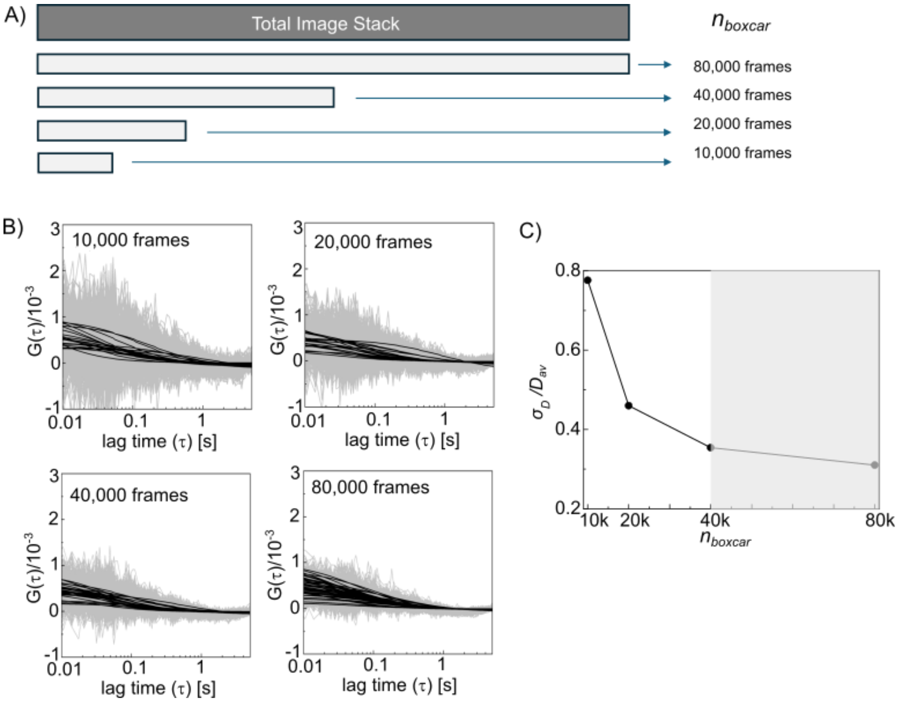


**Figure S3**: Boxcar size ($n_{\text{boxcar}}$) determines quality of ACF fits as evaluated for experimental set-up. A) To optimize $n_{\text{boxcar}}$, an image stack of 80,000 frames was taken from an ROI in resting cells expressing PM-EGFP, and this was divided into smaller sub-stacks of frames (10,000; 20,000; etc.). B) For each sub-stack, ACFs were calculated from all ~625 Px units in the ROI, and these were fit to determine the average and standard deviation of the *D* values ($D_{av}$ and $\sigma_D$, respectively). C) The precision of estimated $D_{av}$, increases with decreasing coefficient of variation ($\sigma_D/D_{av}$). $\sigma_D/D_{av}$ rapidly decreases with increasing frame number until 40,000 frames, after which it changes very little. Based on these results, we selected $n_{\text{boxcar}}$ = 40,000 for our boxcar ImFCS analysis as described in the Main Text.

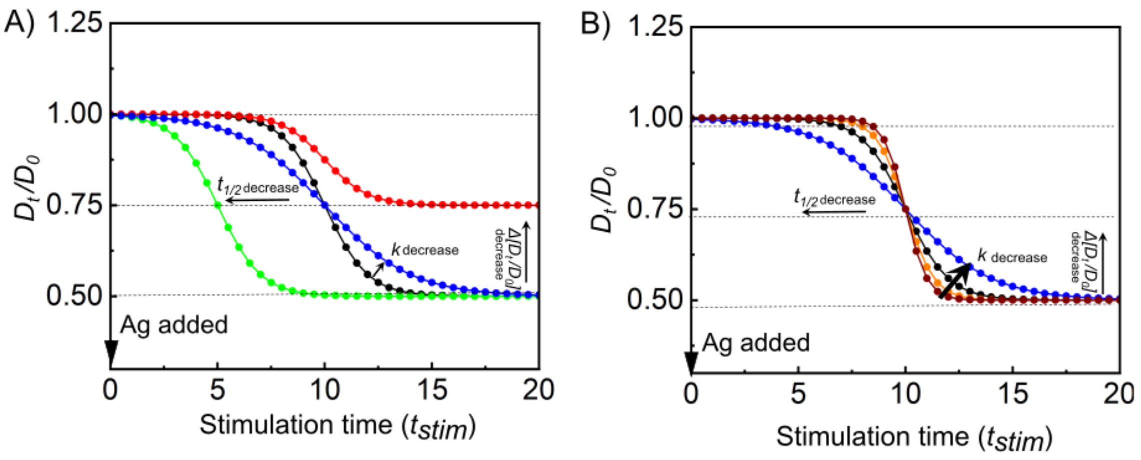


| Parameter | | | | | | |
|---|---|---|---|---|---|---|
| $\Delta[D_t/D_0]$ | 0.5 | 0.25 | 0.5 | 0.5 | 0.5 | 0.5 |
| $k$ | 1 | 1 | 1 | 0.5 | 1.5 | 2.0 |
| $t_{1/2}$ | 10 | 10 | 5 | 10 | 10 | 10 |

**Figure S4**: Parameter values determine curve shape of logistic transition model (Eq 1, Main Text). A) The three adjustable parameters (all with arbitrary units in this illustration) are: $\Delta[D_t/D_0]$, the magnitude of the transition; $t_{1/2}$, the half-time for the transition; $k$, the intrinsic transition rate coefficient. When the absolute value of $\Delta[D_t/D_0]$ decreases, $D_t$ at the stimulated steady state undergoes a smaller change compared to the resting state ($D_0$) (red compared to black). When $t_{1/2}$ is shorter, the entire curve shifts to left (green compared to black). When $k$ decreases, the steepness of the transition curve decreases (blue compared to black). B) For constant $\Delta[D_t/D_0]$ and $t_{1/2}$, the steepness of the transition curve decreases with decreasing $k$ (brown to orange to black to blue).

### Simulation Methods

#### Simulation Setup and Data Acquisition

Simulations were conducted to investigate the lateral diffusion dynamics of identical particles confined within a defined area, using a computational approach analogous to ImFCS measurements. ImFCS analyzes the temporal autocorrelation function (ACF) of fluorescence fluctuations to derive as fitting parameters the diffusion coefficients ($D$) of fluorescent probes.

In the simulation, 51,200 particles were randomly distributed across 180 Px units (18×10 grid), each representing an area of 320×320 $nm^2$. The analysis focused on the central region of 128 Px units (16×8). Particles were initialized with a diffusion coefficient ($D = D_0$) of 0.5 $\mu m^2/s$. Time-dependent changes in diffusion coefficient ($D = D_t$) were modeled using a logistic transition function (Eq. 1, Main Text), parameterized by:

- $\Delta D_{\mathrm{t}}$ (the difference of the *D* values between two steady-states): 0.05 or 0.45 $\mu m^2/sec$
- $\Delta[D_t/D_0]$ = 0.1 (10%) or 0.9 (90%) corresponding to 0.05/0.5 or 0.45/0.5, respectively
- Transition rate coefficient ($k$): 0.01, 0.05, or 0.2 $sec^{-1}$
- Half-time ($t_{1/2}$): 469.0 sec

#### Data Generation

To simulate molecular movement, we used the 2-dimensional random walker algorithm. Briefly, displacement vectors (x, y) were generated by Gaussian distribution with $D$ values over short time intervals of 0.875 msec, assuming *D* value change is negligible within these intervals. Spatial coordinates for each particle were recorded for these time intervals, generating 896,000 location vectors over 784 seconds. To allow for system stabilization prior to change initiation, the initial 4,000 displacement vectors were excluded from analysis.

#### Frame Construction

Frames were created by collecting every fourth vector, resulting in a temporal resolution of 4×0.875 msec = 3.5 msec per frame (to match the experimental time resolution; See Methods section above) and a total of 224,000 frames. Each frame recorded the number of particles within each of the 128 Px units in the central region.

#### Autocorrelation Analysis

Temporal ACFs were calculated for each Px unit to quantify fluctuations in numbers of particles. The first 40,000 frames (spanning 140 seconds) were utilized as the initial dataset. The ACF averaged over all 128 Px units during this and each sequential time interval was computed and fitted to the ImFCS fitting model for two-dimensional diffusion (Eq. S1; (8)),

$$G(\tau) = \frac{1}{N_P}\left(\frac{\mathrm{erf}(p(\tau)) + \frac{\left(e^{-(p(\tau))^2} - 1\right)}{\sqrt{\pi}p(\tau)}}{\mathrm{erf}\left(\frac{a}{\omega_0}\right) + \frac{\omega_0}{a\sqrt{\pi}}\left(e^{-\left(\frac{a}{\omega_0}\right)^2} - 1\right)}\right)^2 + G_\infty; \quad p(\tau) = \frac{a}{\sqrt{4D\tau + (\omega_0)^2}} \tag{S1}$$

where $G(\tau)$ is the ACF as a function of lag time ($\tau$), $N_P$ is the number of particles diffusing within a Px unit, $D$ is the lateral diffusion coefficient in the Px unit, $a$ is the length of the Px unit in the object plane (3.2 nm in the simulation), $\omega_0$ is the point spread function (PSF) of the microscope, $G_\infty$ is the convergence value of $G(\tau)$ at very large lag times. The simulation utilized asymptotic behavior (e.g., for PSF approaching zero in our computational scheme).

Following the initial time interval, ACF calculations were shifted by a specified number of frames (e.g., a 10,000-frame shift corresponding to 35 seconds shift) to capture temporal changes in diffusion dynamics.

**Bootstrap Analysis**

To assess variability in ACF-derived diffusion coefficients, a bootstrapping approach was applied: At each time point, 50% of the ACF curves (64 curves) from individual Px units were randomly selected and averaged. The averaged ACF curves were fitted with the ImFCS equation (Eq. S1) to derive the corresponding time-dependent diffusion coefficients ($D_{\mathrm{t}}$). This process was repeated 30 times, producing a distribution of 30 $D_{\mathrm{t}}$ curves.

**Statistical Analysis**

Each set of $D(t)$ curves was fitted with a logistic function (Eq. 1 in Main Text) to extract parameters, including $k$ (transition rate coefficient) and $t_{1/2}$ (half-time). The 30 replicates of bootstrapped ACFs yield mean and standard deviation of these parameters, allowing subtle changes in diffusion dynamics to be distinguished.

**Boxcar Sizes and Shift**

Two box sizes ($n_{boxcar}$) were used for temporal analysis: A small box comprised 10,000 frames (35 sec). A large box comprised 40,000 frames (140 sec).

Successive ACF calculations were shifted by a set time interval ($\Delta n_{boxcar}$ = 10,000 frames, 35 sec) for each of the two box sizes to capture changes in diffusion dynamics over time.

**Software Implementation**

The simulation and ACF calculations were implemented in MATLAB. Data processing and visualization were performed using MATLAB and Wolfram Mathematica. Bootstrap averaging of ACFs and fitting the averaged ACFs to Eq. S1 were conducted in Mathematica.

**Supplemental Figures (Simulations):**

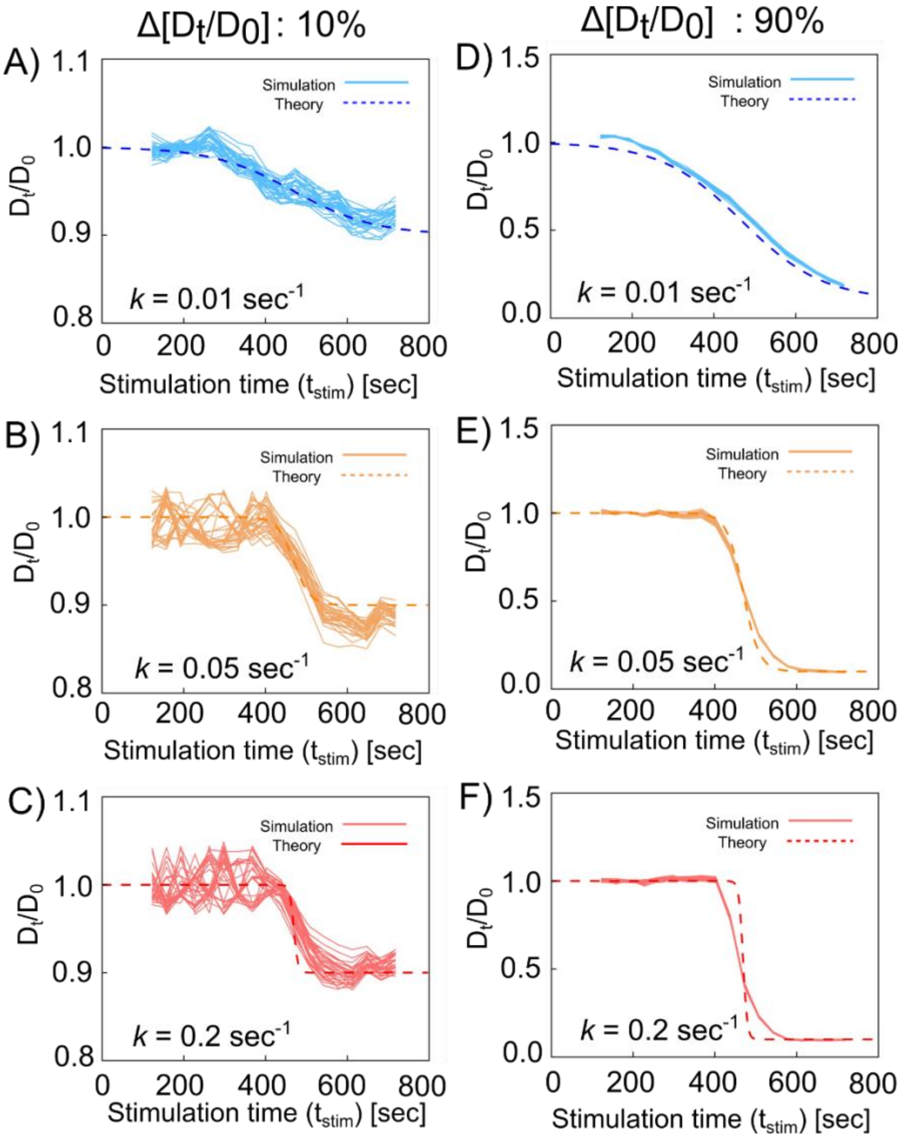


**Figure S5**: Robustness of the logistic model (Eq 1, Main Text) is evaluated by comparing set parameters with simulations. An image stack (128 Px units, 700 sec, 3.5 msec/frame) was simulated using particles with time-dependent changes in diffusion coefficient ($D_t$) determined by Eq. 1 with selected input values for the parameters $\Delta[D_t/D_0]$, $t_{1/2}$ , and $k$. A) Simulations were first carried out with input values like those determined for PM-EGFP: $\Delta[D_t/D_0] = 0.1$, $t_{1/2} = 469$ sec, and $k = 0.01$ sec$^{-1}$ (see Table 2, Main Text). The image stack was processed with overlapping Boxcar ImFCS ($n_{boxcar} = 40{,}000$ and $\Delta n_{boxcar} = 10{,}000$) to obtain $D_t/D_0$ *vs* $t_{stim}$. 30 randomly selected sub-samples (each (64/128 of all simulated Px units) were used to generate respective $D_t/D_0$ plots (solid lines), which can be compared with theoretical curve based on input parameters (dashed line). Averaged values of these parameters from bootstrapped samples agree well with set values as reported in Table S1**.** B-F) For more stringent tests of robustness, additional raw image stacks were simulated using Eq. 1 with increasing $k$ values for $t_{1/2} = 469$ sec and $\Delta[D_t/D_0] = 10\%$ (A-C) or 90% (D-F). Solid lines represent 30 bootstrapped sub-samples and dashed lines show respective theoretical curves. Input values and those extracted from bootstrapped sub-samples are included in Table S1. Comparison shows that simulated data yield quite accurate estimation of $\Delta(D_t/D_0)$ and $t_{1/2}$ in all cases. The $k$ value is accurately estimated for input $k \leq 0.05$ sec$^{-1}$ but significantly underestimated for $k > 0.05$ sec$^{-1}$.

**Table S1: Summary of simulations based on logistic model (Eq. 1 in the Main Text)[a]**

| | **Parameters used for simulation** | | | **Boxcar fitting conditions** | |
|---|---|---|---|---|---|
| | $\Delta\,[D_t/D_0]$ **(%)** | $t_{1/2}$ **[sec]** | $k$ **[sec$^{-1}$]** | $n_{boxcar}$ | $\Delta n_{boxcar}$ |
| Set value | 10% | 469 | 0.01 | 40,000 | 10,000 |
| Fitted value | 9±1% | 427±28 | 0.01±0.01 | | |
| % error | 10% | 9% | 0% | | |
| Set value | 10% | 469 | 0.05 | 40,000 | 10,000 |
| Fitted value | 11±1% | 491±10 | 0.04±0.01 | | |
| % error | 10% | 5% | 20% | | |
| Set value | 10% | 469 | 0.2 | 40,000 | 10,000 |
| Fitted value | 11±0.6% | 476±9 | 0.05±0.007 | | |
| % error | 10% | 2% | 75% | | |
| Set value | 10% | 469 | 0.01 | 10,000 | 10,000 |
| Fitted value | 10±3% | 399±68 | 0.01±0.003 | | |
| % error | 0% | 15% | 0% | | |
| Set value | 10% | 469 | 0.05 | 10,000 | 10,000 |
| Fitted value | 12±3% | 410±131 | 0.04±0.04 | | |
| % error | 20% | 13% | 20% | | |
| Set value | 10% | 469 | 0.2 | 10,000 | 10,000 |
| Fitted value | 10±0.5 | 473±11 | 0.07±0.005 | | |
| % error | 0% | 1% | 65% | | |
| Set value | 90% | 469 | 0.01 | 40,000 | 10,000 |
| Fitted value | 96±11% | 485±34 | 0.01±0.004 | | |
| % error | 7% | 3% | 0% | | |
| Set value | 90% | 469 | 0.05 | 40,000 | 10,000 |
| Fitted value | 90±5% | 471±21 | 0.04±0.002 | | |
| % error | 0% | 0.4% | 20% | | |
| Set value | 90% | 469 | 0.2 | 40,000 | 10,000 |
| Fitted value | 91±6% | 460±7 | 0.05±0.004 | | |
| % error | 1% | 2% | 75% | | |

[a] An image stack (128 Px units, 700 sec with 3.5 msec/frame frame rate) was simulated using particles with time-dependent changes in diffusion coefficient ($D_t$) determined by Eq. 1 (Main Text) using specified values for the parameters $\Delta[D_t/D_0]$, $t_{1/2}$ , and $k$. The image stack was processed with Boxcar ImFCS ($n_{\text{boxcar}}$ = 40,000 or 10,000 and $\Delta n_{\text{boxcar}}$ = 10,000) to obtain $D_t/D_0$. 30 randomly selected sub-samples (each 64/128 of all simulated Px units) were bootstrapped to generate respective $D_t/D_0$ plots. These simulated curves were fit with Eq. 1 to extract parameters that can be compared with set parameters to calculate %error:

$$\%\ error = \left(\frac{Fitted\ value - set\ value}{set\ value}\right) \times 100$$

Values shown are averages and standard deviations of the respective 30 bootstrapped sub-samples.
